# Arp2/3 complex-dependent actin regulation protects the survival of tissue-resident mast cells

**DOI:** 10.1101/2024.02.23.581763

**Authors:** Lukas Kaltenbach, Michael Mihlan, Svenja Ulferts, Mathias Müsken, Katharina M. Glaser, Gerhard Mittler, Magda Babina, Metello Innocenti, Robert Grosse, Theresia E.B. Stradal, Tim Lämmermann

**Author notes:** Correspondence to: Tim Lämmermann.

## Abstract

Actin network dynamics are pivotal in governing the motility and effector functions of immune cells. The Arp2/3 complex is a key regulator of actin filament branching, with mutations in its subunits being linked with human immunodeficiencies. While known for its role in phagocytosis and cell migration, our study uncovers a critical role of the Arp2/3 complex in safeguarding the tissue residency of mast cells (MCs), essential immune cells in allergies, venom detoxification and antigen-specific avoidance. Mechanistically, we show that MCs require Arp2/3-regulated actin filament assembly to resist their integrin-mediated mechano-coupling with their tissue niche. Arp2/3 complex depletion directs MCs into cell cycle arrest and death, which can be rescued by inhibiting their mechanical interactions with extracellular matrix. Our findings underscore the Arp2/3 complex as a mechano-protective element for maintaining MC survival and longevity in tissues, highlighting the importance of actin regulation in preserving the homeostasis of a tissue-resident immune cell population.

**One Sentence Summary:** Arp2/3 complex protects the tissue homeostasis of resident mast cell networks

## INTRODUCTION

Actin cytoskeleton remodeling drives the movement of immune cells, their interactions with other cells and the elimination of microbes. A steadily increasing number of rare primary immunodeficiencies arising from deleterious mutations in genes encoding actin-regulating proteins have been identified over recent years (*1, 2*). Actin nucleators and actin-binding proteins make up the machinery that governs the organization of actin filament networks, determining shape and dynamics of immune cells (*3, 4*). Most of our knowledge on the physiological roles of actin regulation in the immune system comes from immune cell subsets that traffic between organs through the blood and lymphatic system, including T cells, B cells, neutrophils, dendritic cells (DCs) and natural killer (NK) cells. Studies involving human patient material or mouse cells have elucidated critical functions of actin regulation during the adhesion of heavily trafficking immune cells to blood vessels, transmigration through the endothelium, movement through the interstitial tissue space, and interactions with pathogens or other immune cells (*5*). However, the contribution of actin regulation to the homeostasis and tissue residency of sessile immune cells remains poorly understood.

All tissues harbor resident immune cells, many of which capable of self-maintenance over extended time periods. Current concepts on long-term self-maintenance emphasize interactions between resident immune cells and stromal cells of the tissue microenvironment, which create “tissular niches” that mutually benefit both cell types (*6*). These niches provide sessile immune cells a physical foundation to localize in the tissue and factors that regulate their development, survival and tissue-specific imprinting. While many of the critical growth and trophic factors for the tissue-residency of specific immune cell subsets have been identified, the impact of controlling physical interactions with the extracellular milieu on the long-term survival of resident immune cells in the tissue remains largely unclear.

We recently showed that mast cells (MCs), tissue-resident immune cells with important roles during allergic inflammation, anaphylaxis, venom detoxification, and food avoidance (*7–11*), critically rely on adhesive interactions with the extracellular matrix (ECM) to organize their tissue homeostasis and distribution (*12*). MCs utilize integrin receptors, the major family of adhesion receptors in mammals, to anchor themselves to extracellular components in connective tissue. Integrin-mediated adhesions generate cellular sites where actin polymerization and actomyosin contraction transmit intracellular mechanical forces to the surrounding ECM (*13*). In consequence, reciprocal force exchange can occur across integrin-based adhesion between the ECM and the cell. The strong mechanical coupling of MCs with ECM is unique among immune cells, standing in stark contrast to the low-adhesive nature and integrin-independent amoeboid migration of the above-mentioned heavily trafficking immune cell types (*14–16*). Given the formation of prominent integrin-mediated adhesion sites, which connect intracellular actin filaments with the ECM, MCs may have adapted actin-network properties to withstand developing mechanical forces in the tissue. However, the role of actin regulation in maintaining the homeostasis and longevity of tissue-resident MCs has not been explored.

Single cell RNA sequencing data hinted at actin filament branching as a potentially crucial cytoskeletal mechanism for adhesive MCs (*12*). Comparison of WT and adhesion-deficient skin MCs revealed an upregulated transcript expression of *Arpc2*, a component of the seven-subunit actin-related protein 2/3 (Arp2/3) complex (*17*). This complex acts as a dominant nucleator of branched actin networks, binding to the sides of existing actin filaments before initiating the growth of a new filament at a characteristic 70° angle (*18*). Before exerting its actin-nucleating function, the Arp2/3 complex requires activation by nucleation-promoting factors of the WASP and WAVE family proteins (*19, 20*). Branched actin networks, consisting of short and cross-linked actin filaments, support important cellular functions, including the formation of the actin cortex to maintain cellular shape and the generation of pushing forces at the leading edge of crawling cells (*21, 22*). Previous work on immune cells has predominantly focused on the role of the Arp2/3 complex in regulating cell motility and immune cell effector functions. Genetic depletion of individual Arp subunits or pharmacological inhibition of the Arp2/3 complex altered the migration behavior of T cells, DCs, neutrophils, and macrophages (*23–31*). Recent findings have also demonstrated the involvement of the Arp2/3 complex in nuclear actin filament polymerization in T cells (*32*).

Our investigation of the Arp2/3 complex in MCs reveals the essential role of branched actin networks in preserving the longevity of MCs in physiological tissues. We uncover a previously unknown function of the Arp2/3 complex for the survival of MCs in their native environment, highlighting the critical role of actin regulation in maintaining the homeostasis of tissue-resident immune cells that strongly interact with their tissue niches.

## RESULTS

### Arp2/3 complex controls MC morphology and motility in 3D environments

To understand the role of actin filament branching for MC biology, we aimed to deplete the major actin nucleator, the heptameric Arp2/3 complex. Of the seven subunits, ARP2 and ARP3 directly interact with actin filaments, while ARPC2 and ARPC4 form the structural core of the complex (*17*). Therefore, we targeted the *Arpc4* gene to deplete the entire Arp2/3 complex in mouse MCs (*33, 34*). We used bone marrow-derived MCs (BMMCs) that were differentiated toward a connective tissue-type MC (CTMC) phenotype from *Mcpt5-Cre^+/−^ Arpc4^fl/fl^ (Arpc4^ΔMC^)* and control mice (Fig. 1A). Mcpt5-Cre transgenic mice express Cre recombinase in the CTMC compartment, enabling efficient conditional gene targeting (*35*). BMMCs from *Arpc4^ΔMC^* mice showed efficient ARPC4 depletion, leading to the downregulated expression of other Arp2/3 subunits as previously described (figs. S1, A to C) (*33, 34*). Unaltered expression profiles of the cell surface markers c-KIT and FCER1 indicated normal MC differentiation of *Arpc4*-deficient BMMCs (fig. S1D). However, scanning electron microscopy revealed morphological differences between wild-type (WT) and *Arpc4*-deficient cells. WT BMMCs displayed homogenous surfaces with sheet-like extensions, whereas *Arpc4^−/−^* BMMCs showed ruffled extensions with finger-like protrusions (Fig. 1B). When cells were embedded in 3D Matrigel, an artificial gel system that closely mimics MC dynamics of physiological tissues, clear differences in cell morphologies and migration behavior became apparent comparing control and knockout MCs (Figs. 1, C to G, and figs. S1, E and F). The majority of *Arpc4^−/−^* BMMCs displayed elongated and less circular cell bodies with needle-like protrusions at the leading cell edges in comparison to WT BMMCs (Fig. 1C, and figs. S1, E and F). When BMMCs in 3D Matrigel were imaged over 36 h, *Arpc4^−/−^* cells moved faster and covered longer distances than control cells (Figs. 1, D and E, and movie S1). To visualize filamentous (F-)actin organization of 3D migrating cells in more detail, BMMCs from control and *Arpc4^ΔMC^* mice expressing a transgenic Lifeact-GFP reporter were used (*36*). Using spinning-disk confocal microscopy of BMMCs in 3D Matrigel, we confirmed our previous findings that WT MCs form mesenchymal-like shapes with F-actin rich, small lamellopodial-like extensions at the leading edge (Fig. 1F). Strikingly, ARPC4 depletion in MCs caused a very uncommon immune cell morphology. In contrast to WT cells, *Arpc4^−/−^* MCs formed cylindrical cell protrusions covered with bleb-like structures (Fig. 1G, and movie S2). Live cell imaging could visualize that these blebs were occasionally shed as actin-containing fragments from the cell edges or cell bodies of *Arpc4^−/−^* MCs (Fig. 1H, fig. S1G and movie S3). Overall, the cylindrical cell extensions were most similar to previously reported lobopodial protrusions, the formation of which depends on the build-up of intracellular pressure by contractile forces to support the integrin-dependent 3D migration of fibroblasts (*37*). In agreement with this, we found that *Arpc4^−/−^* MCs adopted round morphologies (Figs. 1, I and J) and showed stalled movement in 3D Matrigel (Fig. 1K, and fig. S1H) upon blockade of actomyosin contraction with the Rho kinase inhibitor Y-27632 or the myosin II inhibitor blebbistatin. In contrast, WT MCs retained residual migration speeds, when they migrated as elongated cells in the absence of contractile forces (Fig. 1K, and figs. S1, H to J). Similar to WT cells (Fig. 1M, and figs. S1, K and L), *Arpc4^−/−^* MCs rounded up and halted migration upon blockade of integrin β1 engagement (Figs. 1, L and M, and fig. S1K), confirming the strict dependence of MC movement on integrin-dependent ECM anchoring.

**Fig. 1:**
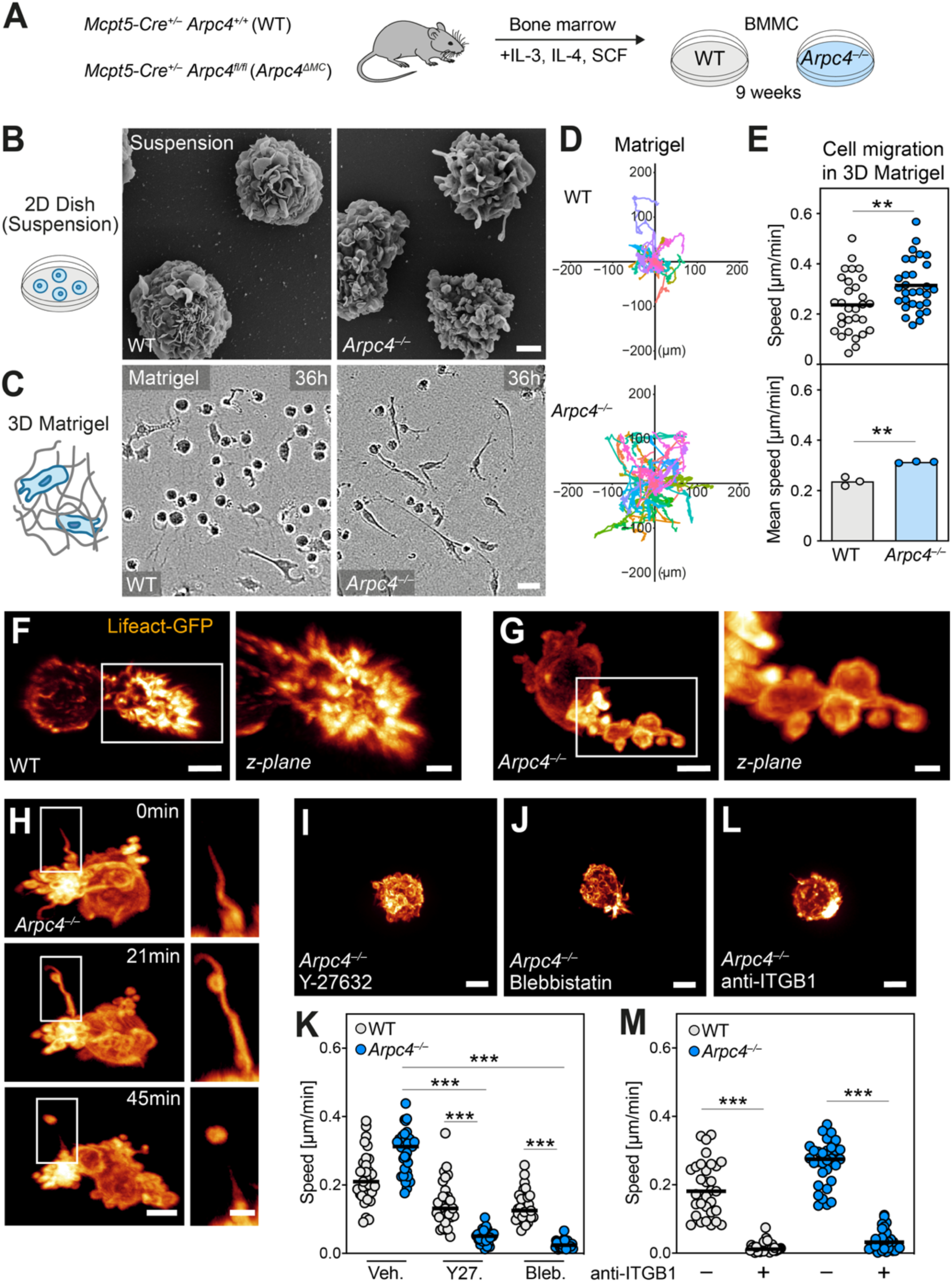
Arp2/3 complex controls MC morphology and motility in 3D environments. **(A)** Scheme for the generation of control (WT) and *Arpc4*-deficient (*Arpc4^−/−^*) bone marrow-derived mast cells (BMMCs) from mice. **(B)** Representative scanning electron microscopy images of WT and *Arpc4^−/−^* BMMCs. **(C– E)** Comparative analysis of WT and *Arpc4^−/−^* BMMC migration in 3D Matrigel. (C) Brightfield images show cell morphologies (*t*=36 h). (D) Trajectories of individual WT and *Arpc4^−/−^* BMMC tracks over 36 h. (E) Quantification of average cell speeds. Dots in the upper graph are values of individuals cells (*N*=30 randomly chosen cells per genotype). Dots in the lower graph show the average cell speed of all cells (*n*=3 independent experiments from 2 biological replicates). Bars displays the mean; \*\**P*≤0.01, *t* test. **(F, G)** Spinning-disk confocal microscopy of Lifeact-GFP (glow) expressing WT and *Arpc4^−/−^* BMMCs migrating in Matrigel. Zoom-ins show an exemplary lamellopodia-like protrusion (WT, F) and lobopodia-like protrusion of the (*Arpc4^−/−^*, G), focal z-plane. **(H)** Spinning-disk confocal microscopy showing the loss of a cellular fragment from the cell edge of a Lifeact-GFP expressing *Arpc4^−/−^* BMMC in 3D Matrigel. A time sequence from live cell imaging over 45 min is displayed. **(I– M)** Testing the lobopodia-like nature of *Arpc4^−/−^* BMMC protrusions in 3D gels. (I, J, L) Spinning-disk confocal images show representative morphologies of Lifeact-GFP expressing cells in 3D gels. To interfere with actomyosin contraction, *Arpc4^−/−^* BMMCs were either treated with Y-27632 (I) or blebbistatin (J). To interfere with integrin β1 (ITGB1)-mediated adhesion, *Arpc4^−/−^* BMMCs were treated with anti-ITGB1 antibody (L). (K, M) Comparative analysis of WT and *Arpc4^−/−^* BMMC migration in fibrillar Matrigel upon functional interference. Quantification of the average cell speed is shown. Dots are values of individual cells (*N*=30 randomly chosen cells per condition). Bars display the median; \*\*\**P*≤0.0001, Dunńs multiple comparison (posthoc Kruskal-Wallis test). Scale bars: 2 µm (B), 50 µm (C), 4 µm, 2 µm (zoom-in) (F, G), 3 µm, 2 µm (zoom-in) (H), 15 µm (I, J, L).

In summary, the absence of Arp2/3-dependent actin filament branching led to substantial adaptations in the MC actin cytoskeleton, affecting MC morphology and motility over 36 h in 3D *in vitro* environments.

### Arp2/3 complex is critical for MC homeostasis in the skin of adult mice

To understand the role of Arp2/3-mediated actin regulation for resident MCs in real tissues, we studied the homeostatic ear skin of mice. Previous work has shown that embryonic skin hosts MCs of yolk-sac origin, which are gradually replaced by MCs originating from definitive hematopoiesis (*38*). By six weeks of age, the dermal connective tissue is entirely populated by definitive MCs (*39*). These long-lived MCs, with slow proliferation rates, establish tissue residency by maintaining themselves independently from the bone marrow (*38*). To assess tissue homeostasis of endogenous skin MCs, we analyzed ear skin whole mount preparations of *Arpc4^ΔMC^* and littermate control mice at different ages (Fig. 2A). Fluorescence-conjugated avidin, which specifically binds to the intracellular MC granules, was utilized to detect MCs in tissues (*40*). Comparison of dermal MC numbers and distribution in these animals revealed an unexpected phenotype. While WT mice maintained a stable MC network over 27 weeks, *Arpc4^ΔMC^* mice gradually lost the dermal MC population with age, showing a substantial drop in tissue-resident MC numbers by 27 weeks (Figs. 2, B and C). Although MCs nearly disappeared in some tissue regions, the overall composition and geometry of stromal tissue elements remained unaffected (fig. S2A). Thus, Arp2/3-dependent actin regulation is critical for maintaining the tissue-resident MC population in adult skin. This highlights a major physiological role of the Arp2/3 complex in preserving MC homeostasis, demonstrating that actin regulation safeguards self-maintaining resident MCs.

**Fig. 2:**
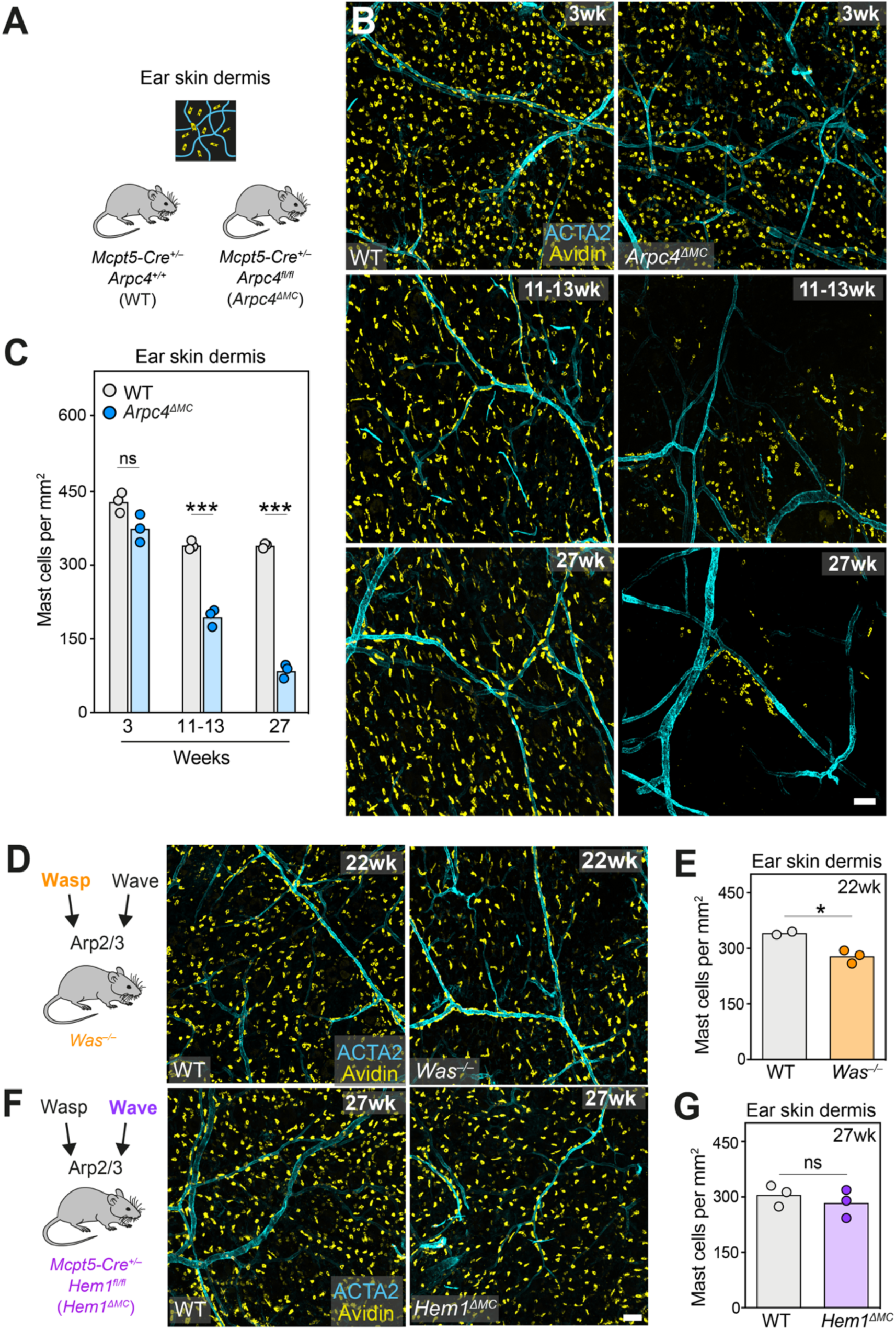
Arp2/3 complex is critical for MC homeostasis in the skin of adult mice. **(A, B)** Representative overview of MC distribution in the ear skin dermis of different mouse age groups. Immunofluorescence stainings of ear skin whole mount tissues from *Arpc4^ΔMC^* and littermate controls. Dermal MCs were immuno-stained with fluorescent avidin (yellow). Arterioles and venules were stained against α-smooth muscle actin (ACTA2, cyan). **(C)** Histological quantification of MC density in the ear skin dermis of *Arpc4^ΔMC^* and littermate control mice at different ages. Each dot represents the average value of three imaging field of views from one mouse (*n*=3 mice per genotype). Bars display the mean; ns, non-significant, \*\*\**P*≤0.0001, *t* test. **(D–G)** The functional contribution of the Arp2/3 upstream activators WASP and HEM1 for MC distribution in the ear skin dermis of adult mice was analyzed. Immunostainings and quantifications were performed as in B and C, respectively. (E, G) Each dot represents the average value of three imaging field of views from one mouse (*n*=2-3 mice per genotype). Bars display the mean; \**P*≤0.05 (*Was^−/−^*), ns, non-significant (*Hem1^ΔMC^), t* test. Scale bars: 100 µm (B, D, F).

Next, we aimed to identify whether a particular Arp2/3 upstream activator controls Arp2/3 complex function in MCs. Therefore, we analyzed mice with genetic deficiencies for the Wiskott-Aldrich Syndrome protein (WASP) and the WAVE complex component hematopoietic protein 1 (HEM1), two proteins shown to play functional roles in other myeloid immune cells (*41–43*). We analyzed ear skin whole mount tissues of constitutive WASP-deficient mice (*Was^−/−^*) and mice with CTMC-specific HEM1-deficiency (*Mcpt5-Cre Hem1^fl/fl^*, short: *Hem1^ΔMC^*). Surprisingly, dermal MC numbers in ≥22 wks-old mice were only moderately affected in *Was^−/−^* mice, and not altered at all in *Hem1^ΔMC^* mice (Figs. 2, D to G). Thus, depletion of either one of these two proteins could not replicate the strong physiological phenotype of *Arpc4^ΔMC^* mice. In line with minor roles of these proteins in controlling MC actin filament organization, *Was^−/−^* and *Hem1^−/−^* BMMCs showed no differences in their 3D morphologies or migration speed over 36 h in Matrigel (figs. S2, B and C).

### Arp2/3 complex protects MC proliferation and survival in 3D Matrigel, but not in suspension

To investigate how Arp2/3-mediated actin regulation protects the longevity of MCs, we adapted our 3D Matrigel model for live-cell imaging of BMMCs over several days. Cultivating WT cells in this environment led to cell growth and an increase in cell numbers over 7 days (Figs. 3, A and B). In contrast, 3D cultures of *Arpc4^−/−^* cells exhibited a proliferation defect with a significant increase in dying cells over the same time period (Figs. 3, A to C, and movie S4). Importantly, we did not observe any defects in cell growth or death when maintaining *Arpc4^−/−^* BMMCs as regular suspension culture over 7 days (Figs. 3, D and E). Similar results were obtained when treating 3D cultured WT BMMCs with the specific Arp2/3 inhibitor CK666 (figs. S3, A to E). These experiments revealed a context-dependent function of Arp2/3-mediated cell growth and protection from cell death, suggesting that actin filament branching is particularly important for MC survival when cells interact with the extracellular environment.

**Fig. 3:**
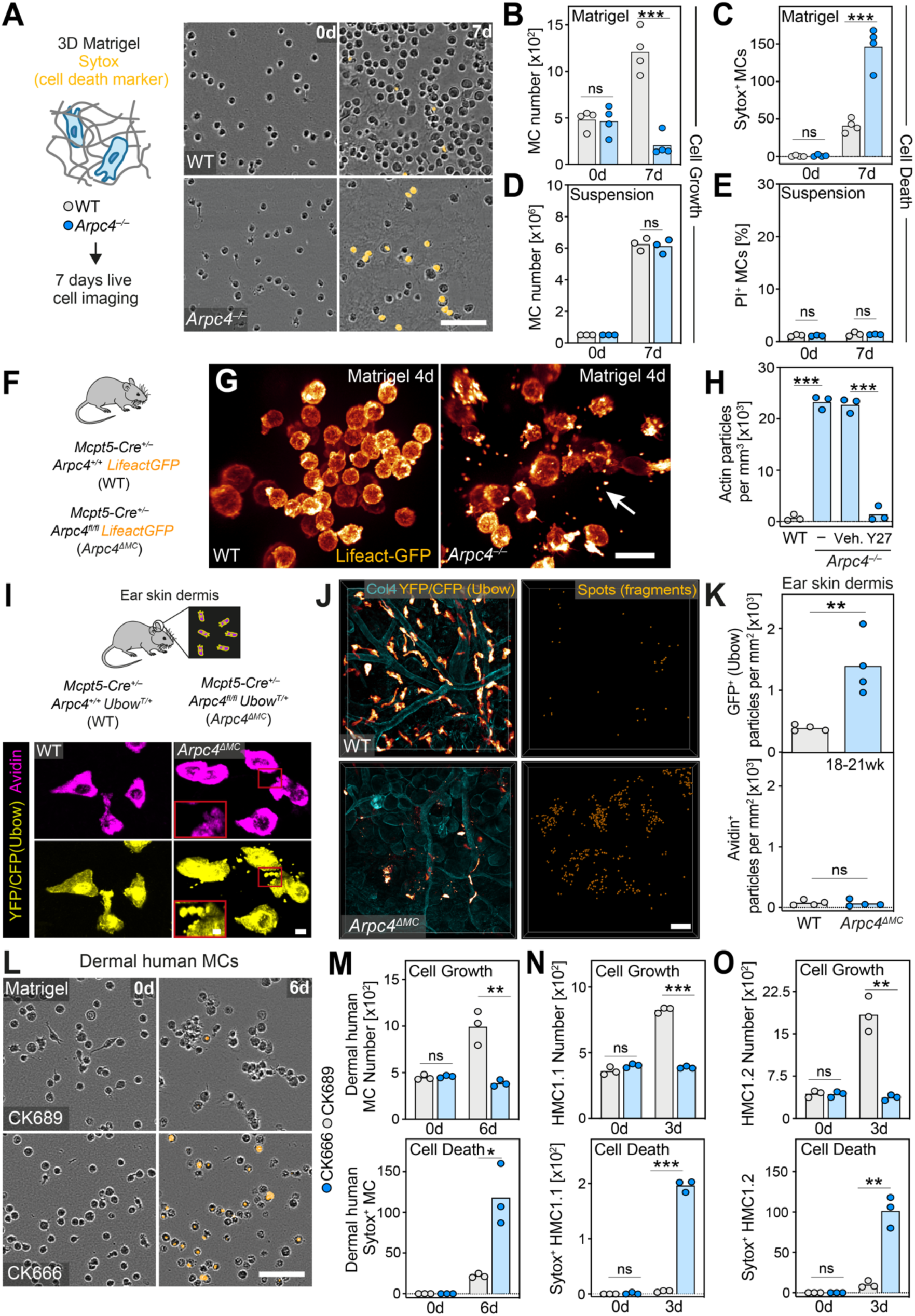
Arp2/3 complex protects MC proliferation and survival in 3D Matrigel, but not in suspension. **(A)** To mimic 3D aspects of the skin dermis *in vitro*, WT and *Arpc4^−/−^* BMMCs were embedded in 3D Matrigel and their behavior monitored over seven days. Brightfield images of WT and *Arpc4^−/−^* MCs in the presence of the cell death marker Sytox (orange) are shown (*t*=0 d, *t*=7 d). **(B–E)** Cell proliferation and cell death of WT and *Arpc4^−/−^* BMMCs was quantified in 3D Matrigel (B, C) or suspension culture (D, E) (*t*=0 d, *t*=7 d). Each dot represents one independent experiment. Bars display the mean. Cell death in the Matrigel was assessed by quantification of Sytox positive BMMCs. Cell death in suspension was quantified by flow cytometric measurements of the percentage of propidium iodide (PI) positive cells. (B, C) *n*=4 independent experiments from 3 biological replicates; ns, non-significant, \*\*\**P*≤0.001; *t* test. (D, E) *n*=3 independent experiments from 2 biological cultures; ns, non-significant, *t* test. **(F)** To visualize the actin cytoskeleton of BMMCs, *Arpc4^ΔMC^* mice were crossed with Lifeact-GFP transgenics. **(G)** Spinning-disk confocal microscopy of Lifeact-GFP expressing *Arpc4^−/−^* BMMCs embedded in 3D Matrigel over several days. Lifeact-GFP positive small actin particles accumulate around cells (highlighted with arrow). **(H)** The number of individual actin particles were measured per defined volume. Y-27632 was applied to interfere with actomyosin contraction. Each dot represents the average value of three imaging fields of view from one independent experiment (*n*=3 independent experiments from 2 biological replicates). The bars display the mean; \*\*\**P*≤0.001, *t* test. **(I)** *Arpc4^ΔMC^* and littermate control mice were crossed with Ubow transgenic mice, which leads to the expression of either YFP or CFP in dermal MCs. Immunofluorescence stainings of ear skin whole mount tissues from WT Ubow and *Arpc4^ΔMC^* Ubow mice are shown. Dermal MCs were immuno-stained with fluorescent avidin (purple) and anti-GFP to amplify YFP and CFP signals. **(J, K)** Histological quantification of avidin- and Ubow-positive fragments in the dermis of ear skin whole mount tissues of WT Ubow and *Arpc4^ΔMC^* Ubow mice. (J) Representative overview images of MCs (orange) in relation to collagen IV (COL4)-expressing basement membrane structures (blue). Fragments are highlighted as spots. (K) Each dot represents the average value of three imaging field of views from one mouse (*n*=4 mice per genotype). Bars display the mean; \*\**P*≤0.01 (Ubow), ns, non-significant, *t* test. **(L, M)** CK666-mediated Arp2/3 complex inhibition on primary human MCs embedded in 3D Matrigel. (L) Representative brightfield images of MCs treated with CK666 or the biological inactive form CK689 in the presence of the cell death marker Sytox (orange) (*t*=0 d, *t*=6 d). (M) Quantification of cell proliferation and cell death. Each dot represents one independent experiment (*n*=3 independent experiments from three different donor pools). Bars display the mean; ns, non-significant, \*\**P*≤0.01 (*t*=6 d, growth), \**P*≤0.05 (*t*=6 d, death), *t* test. **(N, O)** CK666-mediated Arp2/3 complex inhibition on the human mast cell lines HMC1.1 (N) and HMC1.2 (O) embedded in 3D Matrigel. (N) Bars display the mean; \*\*\**P<*0.0001 (*t*=3 d, growth, death), *t* test. (O) Bars display the mean; \*\**P<*0.01 (*t*=3 d, growth, death), *t* test. Scale bars: 80 µm (A), 30 µm (G), 5 µm, 2 µm (zoom-in) (I), 50 µm (J), 80 µm (L).

Since we had observed that *Arpc4^−/−^* BMMCs lost bleb-like cellular fragments in 3D short-term cultures (Fig. 1H), we hypothesized that this phenomenon could contribute to the cell death phenotype in long-term cultures. To address this, we visualized the F-actin organization of BMMCs generated from Lifeact-GFP expressing control and *Arpc4^ΔMC^* mice (Fig. 3F) after embedding them for 4 days in 3D Matrigel. Indeed, substantial increases of F-actin-containing cell fragments occurred around *Arpc4^−/−^* cells, but not for WT BMMCs (Figs. 3, G and H). Y-27632-mediated inhibition of actomyosin contraction completely blocked this phenotype, showing that MC contractility causes fragment formation in the absence of Arp2/3-controlled actin regulation (Fig. 3H). To examine whether such MC fragments also occurred in real tissues, we refined our *Arpc4^ΔMC^*mouse model. While MC detection by fluorescent avidin staining provides strong signals for granule-rich MC bodies, it leaves cellular extensions often weakly stained. By crossing *Arpc4^ΔMC^* mice to Ubow reporter transgenics (*44*), we introduced a cell-specific, cytoplasmic expression of either cyan or yellow fluorescent protein (CFP or YFP) into *Arpc4^−/−^* CTMCs, which allowed the visualization of both cell bodies and protrusions (Fig. 3I). Careful examination of ear skin whole mount tissues revealed CFP/YFP-positive cell fragments around dermal MCs in *Arpc4^ΔMC^* mice, which were only rarely visible in control mice (Figs. 3, I to K). Since fragments were only positive for cytoplasmic CFP/YFP and not stained by the granule marker avidin (Figs. 3I and K, and fig. S3F), we could rule out that MC-released granules contributed to the shed cell fragments. Occasionally, we even detected dermal MCs with blebby lobopodia-like cell extensions, in agreement with our observations *in vitro* (Fig. 3I, zoom-in). Together, our results show that Arp2/3-mediated actin filament organization controls the cellular integrity of the MC actin cortex *in vitro* and *in vivo*.

Several forms of cell-fragment shedding have previously been reported for migrating immune cells (*45–48*). Hence, we wondered whether fragments released from *Arpc4^−/−^* BMMCs related to any of those structures. To address this, we established protocols to purify MC fragments from *Arpc4^−/−^* cells and analyze their composition with mass spectrometry (figs. S3, G to I). Functional annotations and top fifty protein hits revealed a very broad protein composition, which we could not relate to any known shed cell structure. In addition, we could not establish a clear connection between the detected proteins in MC fragments and the observed proliferation defect of *Arpc4^−/−^* BMMCs (figs. S3, J and K). However, we ruled out that the bleb-like fragments of *Arpc4^−/−^* MCs related to apoptotic blebs. When Matrigel-embedded MCs were starved to induce MC apoptosis, apoptotic blebs were not released and remained attached to cell bodies (fig. S3L). Thus, Arp2/3 complex depletion causes the release of MC fragments, which appear unrelated to previously described migration-induced cell fragments, such as migrasomes (*45*). Moreover, we addressed whether the cell death phenotype of *Arpc4^−/−^* BMMCs was related to the process of cytothripsis, *i.e.*, migration-induced cell shattering, which had previously been described for T cells and DCs (*47, 48*). Tracking analysis confirmed our earlier results, showing that *Arpc4^−/−^* BMMCs were more migratory than WT cells (Fig. 1E), which also reflected in a higher fraction of moving cells, before they underwent cell death (fig. S4A). However, we found only anecdotal evidence that migratory *Arpc4^−/−^* BMMCs underwent shattering into pieces, which then resulted in cell death (fig. S4B). The majority of dying *Arpc4^−/−^* cells stopped movement as intact cells shortly before losing their membrane integrity (movie S4). Hence, the Arp2/3 complex prevents cell death, which appears unrelated to cytothripsis.

Given our findings on mouse MCs, we investigated whether Arp2/3-mediated actin regulation similarly protects the survival of human MCs in 3D environments. Therefore, we used (i) primary MCs from human foreskin tissue (Figs. 3, L and M), and (ii) two human MC lines, which had initially been established from a patient with MC leukemia (Figs. 3N and O, and figs. S4, C and D), studying their behavior in 3D Matrigel. In agreement with our results on mouse MCs, human MCs displayed similar growth defects and substantial cell death in response to CK666-induced Arp2/3 inhibition, which did not occur upon treatment with CK689, the biologically inactive form of CK666 (Figs. 3, L to O, and figs. S4, C and D).

In summary, we unveil a previously unrecognized role of the Arp2/3 complex in immune cells, emphasizing the essential role of Arp2/3-mediated actin filament branching in protecting the survival of mouse and human MCs, when cells anchor to surrounding 3D ECM.

### MCs require the Arp2/3 complex for cell cycle control in 3D Matrigel

To gain a more detailed understanding of the proliferation defect observed in *Arpc4^−/−^* MCs, we investigated the nuclear shapes of 3D-matrix embedded cells. Intriguingly, the majority of *Arpc4^−/−^* BMMCs exhibited abnormal nuclear morphologies indicative of incomplete cell divisions, which was scarcely observed in control cells (figs. S5, A and B). This prompted us to explore the involvement of the Arp2/3 complex in cell cycle regulation. BMMCs were retrovirally transduced with the nuclear FastFUCCI reporter, distinguishing G1/S (red) and G2/M (green) cell cycle stages, and their behavior in 3D Matrigel imaged over seven days (Figs. 4, A and B, and movie S5). Proliferating control MCs showed increased percentages of dividing cells (Figs. 4, C and D) and increased numbers of cells transitioning between G1/S and G2/M (Figs. 4, E and F, and movie S5). In contrast, *Arpc4^−/−^* BMMCs in 3D cultures were mostly non-dividing, displaying reduced numbers in G1/S and G2/M cell cycle stages (Figs. 4, C to F). Treatment of WT BMMCs with the Arp2/3 complex inhibitor CK666 yielded similar effects on cell division and cell cycle stages, phenocopying the genetic model (figs. S5, C to G). Importantly, no discernable effects on cell cycle stages were observed when *Arpc4^−/−^*and control BMMCs were grown in suspension (Fig. 4G), underscoring the impact of the environmental context on the observed phenotype.

**Fig. 4:**
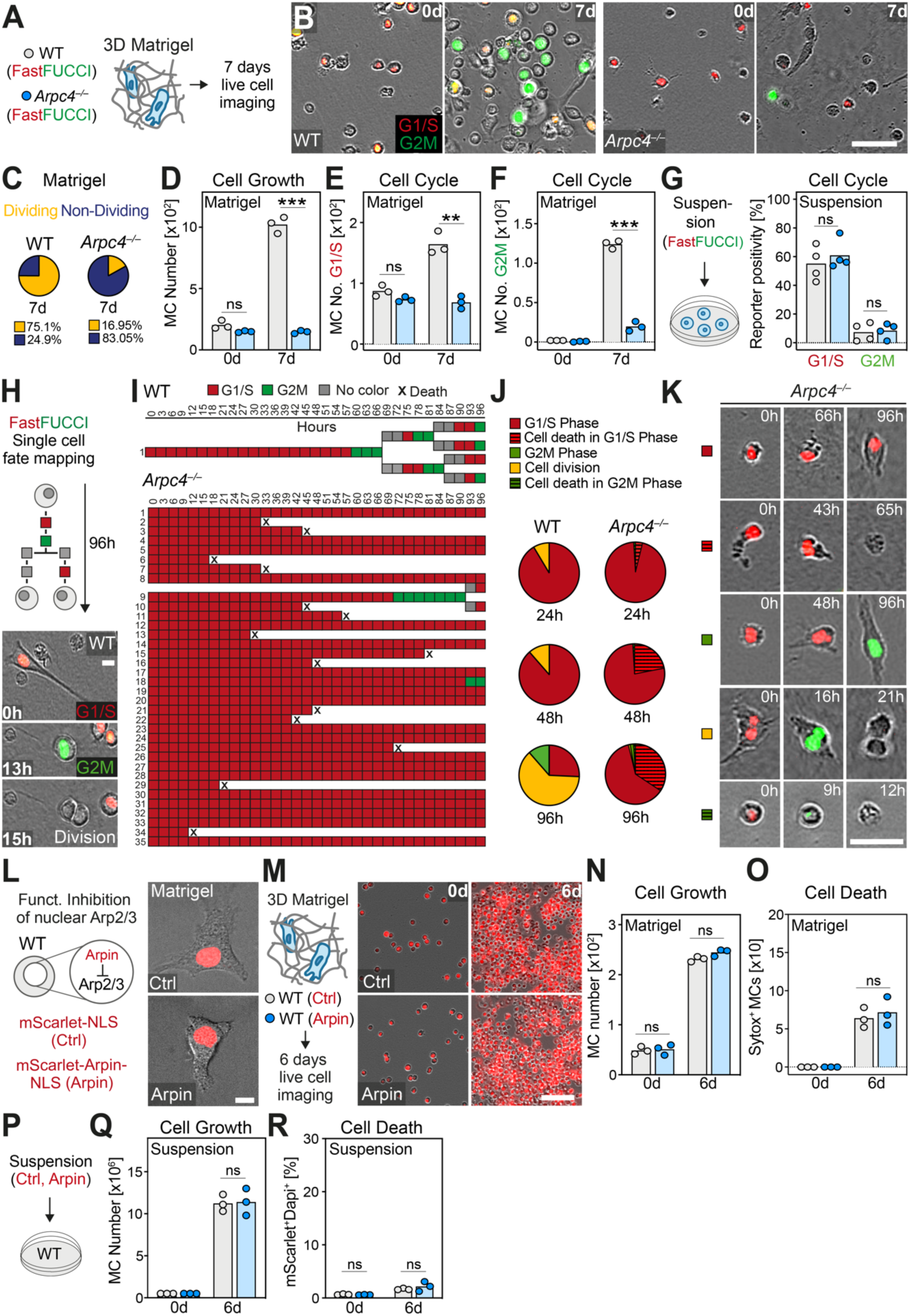
MCs require the Arp2/3 complex for cell cycle control in 3D Matrigel. **(A)** Cell cycle analysis of growing WT and *Arpc4^−/−^* BMMCs in fibrillar 3D gels. WT and *Arpc4^−/−^* BMMCs were transduced with FastFUCCI to differentiate cell cycle stages by nucleus color: G1/S phase (red), G2/M phase (green). **(B–F)** FastFUCCI transduced WT and *Arpc4^−/−^* BMMCs were monitored in 3D Matrigel over seven days. (B) Representative brightfield and epifluorescence images are shown (*t*=0 d, *t=*7 d). (C) Quantification of dividing and non-diving MCs over seven days. Percentages represent average values obtained from *n*=3 independent experiments per genotype. (D–F) Quantification of cell proliferation and cell cycle stages (*t*=0 d, *t=*7 d). Each dot represents one independent experiments (*n*=3 independent experiments per genotype). Bars display the mean; ns, non-significant, \*\*\**P<*0.0001, \*\**P*≤0.01, *t* test. **(G)** Comparative analysis of cell cycle stages of BMMCs in suspension. Reporter positivity was analyzed with flow cytometry. Each dot represents one independent experiment (*n=*4 independent experiments per genotype). Bars display the mean; ns, non-significant, *t* test. **(H)** Single cell fate mapping analysis based on microscopic data on cell division and cell cycle stages was performed over 96 h for WT and *Arpc4^−/−^* BMMCs in 3D Matrigel. **(I)** Visualization of cell cycle dynamics of WT and *Arpc4^−/−^* BMMCs in 3D Matrigel by single cell fate mapping every 3 h. Each colored box represents a cell cycle stage, cross indicates cell death. A representative analysis of 1 WT cell and 35 *Arpc4^−/−^* BMMCs is shown. **(J)** Quantification of cell cycle fates over time. For *Arpc4^−/−^* BMMCs the percentage values derive from the mean value of three independent experiments per genotype, for WT BMMCs the percentage values derive from the mean value of one independent experiment. For each independent experiment 35 randomly chosen WT and *Arpc4^−/−^* BMMCs were analyzed over 96 h. **(K)** Representative time-lapse sequences of cell fates of *Arpc4^−/−^* BMMCs over time in 3D gel are shown. **(L)** The contribution of nuclear Arp2/3 on MC proliferation and survival in 3D Matrigel was analyzed. WT BMMCs were transduced with mScarlet-Arpin-NLS to specifically inhibit Arp2/3 in the nucleus. **(M)** Transduced control (Ctrl) and Arpin-expressing BMMCs were embedded in 3D Matrigel and monitored over six days. Representative images are shown (*t*=0 d, *t=6* d). **(N–R)** Cell growth and survival of Ctrl and Arpin BMMCs over 6 days in 3D Matrigel and suspension was analyzed. Each dot represents one independent experiment (*n*=3 independent experiments per genotype). (N, O, Q, R) Bars display the mean; ns, non-significant, *t* test. Scale bars: 80 µm (B), 20 µm (H), 50 µm (K), 80 µm (M).

Next, we performed single cell fate mapping analysis of Matrigel-cultured MCs, following randomly chosen cells over 96 h and determining their cell cycle stage every 3 h (Fig. 4H). Control BMMCs required a lag time of 2 to 2.5 days before entering their full proliferative phase in 3D Matrigel conditions. Most cells then began to transition from G1/S into G2/M phase and continued to divide in the following time period (Fig. 4I and fig. S5H). Strikingly, *Arpc4^−^*/^−^ BMMCs were mostly found in G1/S phase over the whole 96 h (Figs. 4, I and J). Although different cell fates of *Arpc4^−/−^* BMMCs could be categorized (Fig. 4K and movie S5), the majority of knockout cells remained arrested in G1/S phase with a substantial fraction of cells undergoing cell death over extended time periods (Figs. 4, I and J, and movie S5). Thus, Arp2/3-mediated actin filament branching plays a crucial role in controlling the transition into G2/M phase, a critical process for proliferative MCs establishing their population size in a microenvironment that they strongly interact with.

Since recent studies, mostly in non-immune cells, highlighted functional roles of the Arp2/3 complex in the nucleus (*49, 50*), we explored whether nuclear Arp2/3 might contribute to the MC phenotype. To block nuclear Arp2/3 complex function, we employed a previously established experimental strategy involving the expression and shuttling of Arpin protein, a natural inhibitor of the Arp2/3 complex, to the nucleus (*51*). WT BMMCs were retrovirally transduced with mScarlet-Arpin-NLS or control mScarlet-NLS, which both localized to cell nuclei (Fig. 4L). Similar to control BMMCs, Arpin-expressing BMMCs did neither show a proliferation defect nor increased cell death in 3D Matrigel or suspension cultures (Figs. 4, M to R). Thus, our results argue against an involvement of nuclear Arp2/3 in MC growth, survival and cell cycle progression. Instead, our combined findings point toward a critical role of cytoplasmic Arp2/3-mediated actin filament branching in regulating G1/S to G2/M phase transitions of haptokinetic MCs.

### Arp2/3 complex is critical for MC proliferation and survival upon mechano-coupling

Our previous findings consistently indicated a protective function of cytoplasmic Arp2/3 in sustaining the survival of proliferating MCs that mechanically couple to the ECM in 3D Matrigel or *in vivo*. Assessing the adhesion of *Arpc4^−/−^*BMMCs to various ECM components coated on 2D culture dishes, we could not detect any gross changes in comparison to WT cells. Both BMMC genotypes adhered well to fibronectin, laminin and Matrigel, with the exception of collagen I due to the absence of collagen I-binding integrins (Fig. 5A). Notably, *Arpc4^−/−^*BMMC binding to fibronectin, the most adhesive ligand for MCs, increased the rate of dead cells after 48 h, a phenomenon not observed for control cells (Fig. 5B). Hence, even the adhesive interaction with a planar surface led to a mild compromise in the survival of Arp2/3 complex-depleted MCs.

**Fig. 5:**
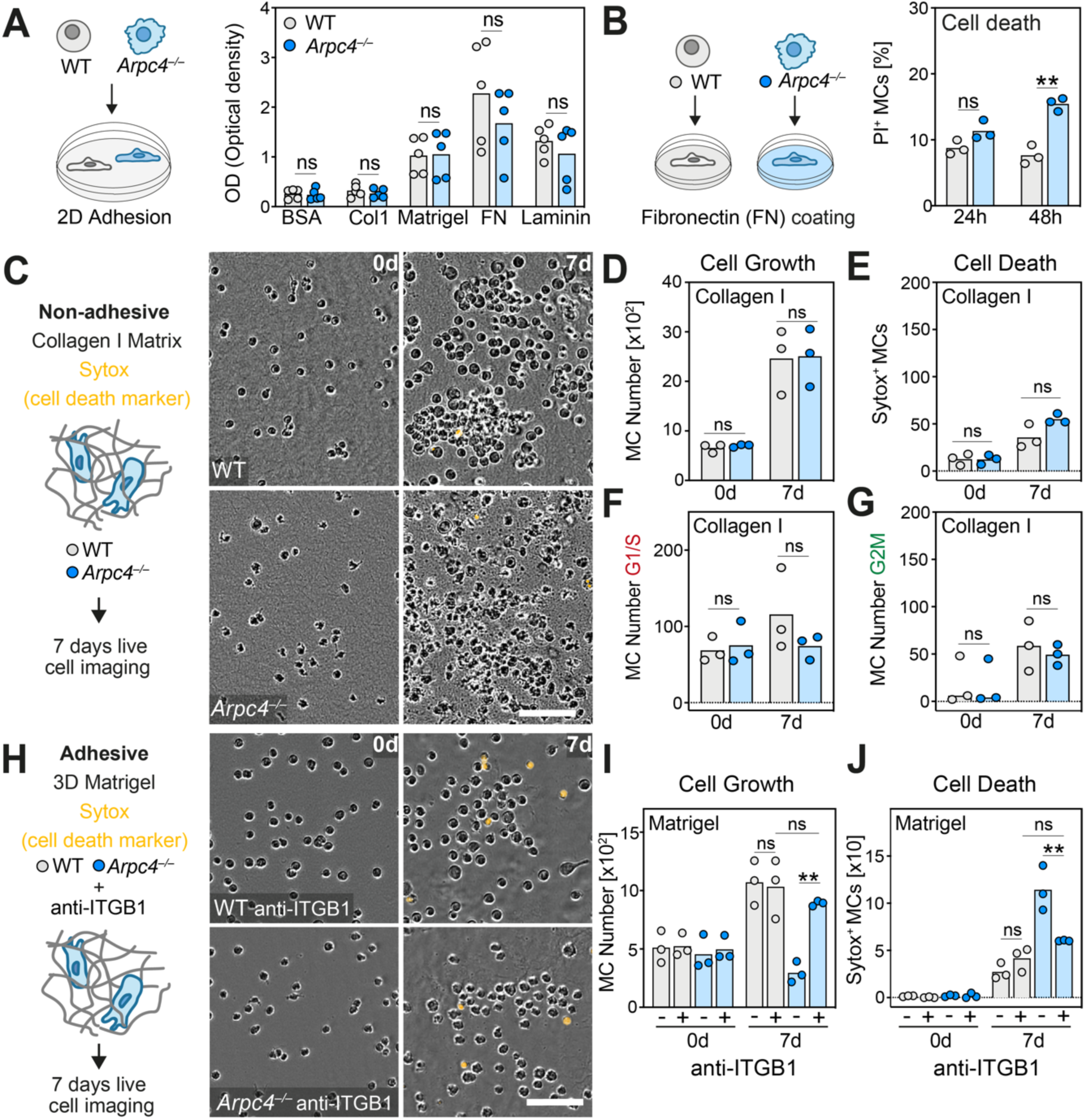
Arp2/3 complex is critical for MC proliferation and survival upon mechano-coupling. **(A)** Adhesion assay of WT and *Arpc4^−/−^* BMMCs to several ligands upon stimulation with KITLG. Bovine serum albumin (BSA) assessed background adhesion. Col1 (collagen 1), FN (fibronectin). Bars display the average optical density (OD) value of adherent cells. Each dot represents an independent experiment (*n*=5 independent experiments from 3 biological replicates) for each condition. Bars display the mean; ns, non-significant; *t* test. **(B)** The contribution of integrin-mediated adhesion on the death of WT and *Arpc4^−/−^* BMMCs under 2D culture conditions was analyzed. WT and *Arpc4^−/−^* BMMCs were kept on FN-treated dishes for 48 h. Flow cytometric quantification of the percentage of PI positive cells at different time points (*t=*24 h, *t*=48 h). Each dot represents one independent experiment (*n*=3 independent experiments from 2 biological replicates). Bars display the mean; ns, non-significant, \*\**P*≤0.01, *t* test. **(C)** WT and *Arpc4^−/−^* BMMCs were embedded in non-adhesive 3D collagen I gels and monitored over seven days. Brightfield images with the cell death marker Sytox (orange) are displayed (*t*=0 d, *t*=7 d). **(D–G)** Quantification of cell proliferation, cell death and cell cycle stages of WT and *Arpc4^−/−^* BMMCs in 3D collagen I gels. Bars display the mean (D–F) or the median (G, 0 d). Each dot represents one independent experiment from *n*=3 independent experiments per genotype. ns, non-significant, *t* test (D–G, 7 d values), *U* test (G, 0 d). **(H)** To rescue *Arpc4^−/−^* BMMCs in adhesive 3D Matrigel, *Arpc4^−/−^* BMMCs were treated with anti-ITGB1 antibody. Brightfield images of WT and *Arpc4^−/−^* BMMCs with anti-ITGB1 treatment and the cell death marker Sytox (orange) are displayed (*t*=0 d, *t*=7 d). **(I, J)** Quantification of cell proliferation and survival of WT and *Arpc4^−/−^* BMMCs with anti-ITGB1 treatment in adhesive 3D Matrigel. Bars indicate the mean; each dot represents one independent experiment (*n=*3 independent experiments per genotype). (I) ns, non-significant, \*\**P*≤0.01, Tukeýs multiple comparison (posthoc one-way Anova test for 7 d values). Scale bars: 80 µm (C, H).

Based on our observations, we hypothesized that the adhesive coupling with the 3D matrix, rather than the 3D geometry itself, induced the death of *Arpc4^−/−^* BMMCs in Matrigel. To test this hypothesis, we disrupted the mechanical coupling of WT and *Arpc4^−/−^* BMMCs in two experimental setups. First, we embedded MCs in non-adhesive 3D collagen I matrices, where both WT and *Arpc4^−/−^* BMMCs adopted round amoeboid-like shapes. Strikingly, *Arpc4^−/−^* MCs proliferated as control cells in 3D collagen gels, without showing any signs of increased cell death over seven days (Figs. 5, C to E). Additionally, cell cycle stages measured by FastFUCCI expression remained unaltered between control and knockout cells (Figs. 5, F and G). Furthermore, *Arpc4^−/−^* BMMCs, expressing Lifeact-GFP, released actin-containing cell fragments in non-adhesive collagen I gels (fig. S6A), akin to what was observed in 3D Matrigel (Figs. 3, G and H). Thus, MCs can proliferate and survive in non-adhesive 3D environments without Arp2/3-controlled actin filament regulation. The formation and release of bleb-like cell fragments in Arp2/3 complex-depleted MCs proved to be an epiphenomenon, which was functionally unrelated to the phenotypic cell cycle arrest and cell death.

Second, we embedded MCs in adhesive 3D Matrigel in the presence of anti-β1 integrin antibody to block integrin-mediated mechanic-coupling with the matrix. Functional integrin blockade switched cellular morphologies to amoeboid-like shapes, confirming the loss of integrin-mediated adhesion over the entire 7 days (Fig. 5H). Under these conditions, both WT and *Arpc4^−/−^*BMMC movement in Matrigel was halted (fig. S6B). Importantly, integrin blockade enabled *Arpc4^−/−^*BMMC to proliferate normally and protected MCs from cell death in Matrigel, rescuing the cellular phenotype of Arp2/3-depleted MCs (Figs. 5, H to J, movie S6). In summary, our study highlights the critical dependence of MCs on Arp2/3-regulated actin networks to ensure proliferation and maintain their survival in the complex ECM. These results underscore that the Arp2/3 complex protects MCs while they adhere and are mechanically coupled to the surrounding environment via integrin receptors.

### Arp2/3 complex protects the tissue residency of mechano-coupled MCs in the skin

Lastly, we aimed to elucidate whether the indispensable role of the Arp2/3 complex in preserving the tissue-resident MC pool within the adult mouse skin is directly tied to mechano-coupling in the physiological tissue. We hypothesized that an additional loss of integrin-mediated adhesion could rescue the depletion of *Arpc4*-deficient MCs in the dermal connective tissue of *Arpc4^ΔMC^*mice. Recently, we investigated mice with MC-specific conditional depletion of talin-1, the key regulator for the conformational switch of integrin receptors from the low-to the high-affinity state, which is required to mediate adhesion to ECM-ligands (*52*). This previous study demonstrated that endogenous MCs of *Tln1^ΔMC^* mice lose their integrin-mediated adhesion in the ear skin dermis, leading to substantial changes in MC shape, migration and tissue positioning. However, MC numbers in the dermis remained unaltered in *Tln1^ΔMC^* mice. Therefore, we crossed *Tln1^fl/fl^* mice with *Mcpt5-Cre^+/−^ Arpc4^fl/fl^* mice to generate *Arpc4/Tln1^ΔMC^* mice harboring MCs with double gene deficiency (Fig. 6A). Comparisons of ear skin whole mount tissues from WT, *Arpc4^ΔMC^*and *Arpc4/Tln1^ΔMC^* littermate mice revealed an astonishing *in vivo* phenotype: the loss of integrin-mediated mechanical coupling allowed *Arpc4*-deficient MCs in *Arpc4/Tln1^ΔMC^* mice to establish a dermal MC pool comparable to that of control mice (Figs. 6, A and B). Further comparison to WT mice showed that *Arpc4/Tln1^ΔMC^*mice displayed slightly altered MC tissue distribution (Fig. 6A), including decreased MC periarteriolar alignment and increased formation of MC clusters in interstitial spaces, which are phenotypes that we had previously reported for *Tln1^ΔMC^* mice. In summary, our *in vivo* data establish Arp2/3-regulated actin filament assembly is essential for the tissue residency of long-living MCs that are mechanically coupled to the ECM. Thus, actin network regulation emerges as a crucial mechanism for maintaining the homeostasis of tissue-resident immune cells, which strongly interact with their tissue niches.

**Fig. 6:**
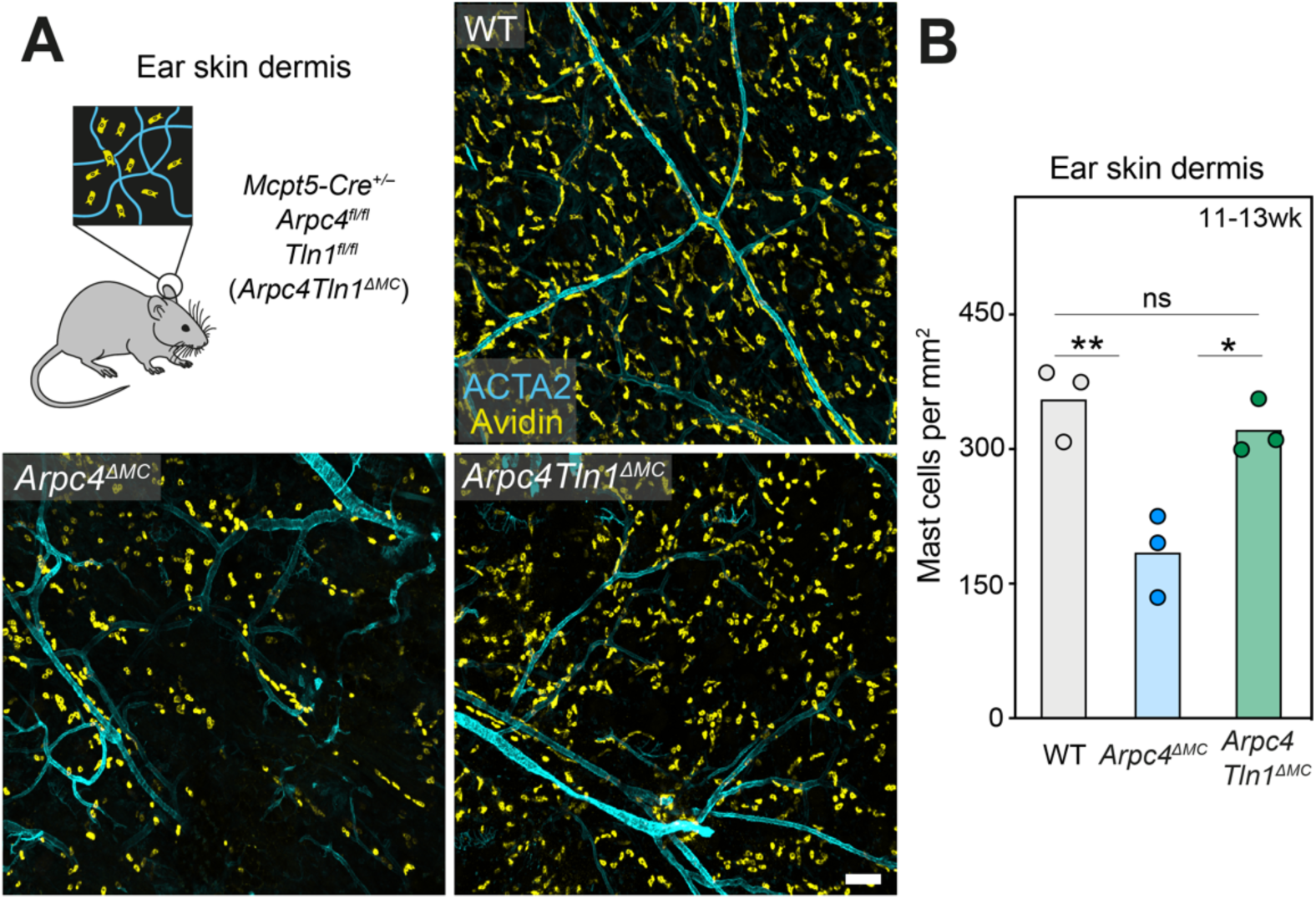
Arp2/3 complex protects the tissue residency of mechano-coupled MCs in the skin. **(A, B)** Testing whether integrin-ECM interactions mediate the *Arpc4^ΔMC^* phenotype of impaired MC homeostasis in native skin tissue. *Arpc4^ΔMC^* mice were crossed with *Tln1^fl/fl^* mice to generate *Mcpt5-Cre^+/−^ Arpc4^fl/fl^Tln1^fl/fl^* (*Arpc4Tln1^ΔMC^)* mice, which have MC-specific functional interference of the Arp2/3 complex and integrin functionality. (A) Representative overview of MC distribution in the ear skin of 11 to 13 weeks old mice. Immunofluorescence stainings of ear skin whole mounts from *Arpc4^ΔMC^*, *Arpc4Tln1^ΔMC^* and littermate control mice. Dermal MCs were immuno-stained with fluorescent avidin (yellow). Arterioles and venules were stained against α-smooth muscle actin (ACTA2, cyan). (B) Histological quantification of MC density in the ear dermis of *Arpc4^ΔMC^*, *Arpc4Tln1^ΔMC^* mice and littermate controls. Each dot represents the average value of three imaging field of views from one mouse (*n*=3 mice per genotype). Bars display the mean; ns, non-significant, \*\**P*≤0.01, \**P*≤0.05, Tukeýs multiple comparison test (posthoc one-way Anova test). Scale bar: 100 µm (A).

## DISCUSSION

Our current understanding of immune cell tissue residency underscores the intricate interactions between resident immune cells and their stromal niche environment. Crucial signaling pathways initiated by growth factors and regulated by transcription factors have been identified as key determinants in establishing and maintaining various immune cell populations within tissues. The development and survival of MCs are particularly contingent on stromal cell-derived KIT ligand, also known as stem cell factor (SCF), which activates the receptor tyrosine kinase KIT (*53, 54*). Additionally, transcription factors GATA binding protein 2 (GATA2) and Microphthalmia-associated transcription factor (MITF) play pivotal roles in MC differentiation. Mouse models deficient in either of these molecules exhibit a complete absence or substantial reduction of MCs in physiological tissues (*53–56*).

In this study, we shed light on the critical role of the actin filament network in governing the long-term maintenance of resident MC populations *in vivo*. Our research identified a previously unrecognized function of Arp2/3-mediated actin network control in immune cell proliferation and cell cycle regulation, a phenomenon not yet reported for other immune cell types. Previous studies on neutrophils, macrophages, T cells, NK cells, and DCs highlighted the roles of Arp2/3-controlled actin network expansion in functions such as phagocytosis (*30, 57, 58*), granule exocytosis (*59*), lamellipodia formation at the cellular leading edge (*24, 26, 28–30*), transendothelial migration (*25, 60*) and space exploration in complex environments (*23, 27, 31, 61, 62*). Further investigations emphasized roles of the Arp2/3 complex for perinuclear actin in immune cell migration (*63*) and nuclear actin in the process of T cell activation (*32*). Genetic defects in individual components of the Arp2/3 complex, with a focus on ARPC1B and ARPC5 isoforms, have been associated with immune system disorders in humans. The reported defects, resulting in combined immunodeficiencies with severe inflammation and early-onset autoimmunity, inflammation, and mortality, highlight the clinical relevance of Arp2/3 complex dysfunction (*64–66*). Notably, isoform diversity within the Arp2/3 complex may determine cell-specific phenotypes (*67, 68*), prompting further exploration of individual ARPC isoforms for MC biology.

Our research emphasizes the pivotal role of Arp2/3-mediated actin filament branching in safeguarding the survival of proliferating MCs, particularly when these cells adhere and are mechanically coupled to the surrounding ECM. Unlike many other immune cell types (*15*), MCs anchor themselves to the tissue niche via β1-integrin receptors, stably connecting intracellular actin filaments with the external environment (*12*). Surprisingly, MCs in the skin dermis or in adhesive Matrigel cannot tolerate the absence of the Arp2/3-regulated actin networks, causing cell cycle arrest and death. However, this phenotype can be rescued by uncoupling MCs from the ECM through integrin blockade. Additionally, MCs exhibit normal proliferation in non-adhesive environments. Therefore, the protective role of the Arp2/3 complex in proliferating MCs appears functionally connected to the cells’ pronounced haptokinetic nature. In line with this notion, similar phenomena related to cell proliferation and death have not been reported for low-adhesive immune cell types lacking Arp2/3 complex function.

Previous studies on non-immune cells have emphasized the critical roles of actin filament assembly in interphase and mitosis (*69*). Our results identified a cell cycle block in G1/S phase in MCs that bind to Matrigel components via integrin receptors in the absence of Arp2/3-mediated actin filament branching. This suggested that perturbed integrin signaling could contribute to the proliferation arrest. In the vast majority of non-immune cells, integrin-mediated cell adhesion transmits signals to promote cell cycle progression through G1 phase, and loss of adhesion causes G1 cell cycle arrest (*70, 71*). However, unlike fibroblasts and other substrate-dependent cells, MCs do not depend on integrin signaling for their proliferation. MC numbers remain consistent in mouse models with MC-specific integrin deficiencies, and MCs exhibiting compromised integrin function proliferate normally in 3D Matrigel cultures (*12*). Therefore, it is unlikely that integrin signaling events play a role in Arp2/3-regulated cell cycle progression.

Previous *in vitro* evidence in fibroblasts suggested that actin filaments might be sensed inside the cell, delivering an essential “go” signal for DNA synthesis and cell growth during the cell cycle (*72, 73*). Disrupting actin filaments by cytochalasin treatment resulted in a G1 cell cycle block, which could be alleviated when p53 or Rb tumor suppressors were inactivated (*74, 75*). Knockdown of the Arp2/3 complex in fibroblasts also halted cell proliferation unless the cyclin-dependent kinase (CDK) inhibitor p16 was inactivated (*76*). Based on these studies, a cell cycle checkpoint that senses branched actin networks and thus informs the cell about correct cytoskeleton dynamics, was speculated to provide essential signals for cell cycle progression (*72*). Refined experiments with inducible depletion of the Arp2/3 complex in fibroblasts confirmed the G1 cell cycle block and emphasized that multiple nuclear and cytoplasmic signaling factors collaborate during the induction and maintenance of the proliferation arrest (*77*). Analysis of Arp2/3-depleted fibroblasts cultivated in conventional 2D in vitro systems highlighted multiple molecular consequences, including abnormal spindle actin and microtubule abnormalities, the formation of micronuclei after incomplete mitosis, DNA damage, p53 activation, CDK inhibitor p21-mediated cell cycle arrest, cGAS/STING signaling, and cellular senescence (*77*). Such detailed cellular and biochemical analyses are challenging for MCs embedded in 3D Matrigel. However, we hardly observed Arp2/3-depleted MCs undergoing mitosis in live-cell imaging experiments over several days and found no evidence for the formation of micronuclei or DNA damage (data not shown).

In addition to biochemical signals, mechanical forces have emerged as crucial regulators of the G1-S phase transition (*70, 78*). Since branched actin filament networks control the cell shape and cytoskeletal organization, the absence of the Arp2/3 complex might predominantly alter the biophysical force balance inside MCs. Studies in adhesive non-immune cells have demonstrated that intracellular tension controlled by actomyosin contractility is linked to cell cycle progression through G1/S phase (*79–81*). Due to ECM coupling, mechanical forces from the environment could readily transmit to MCs, which may experience additional mechanical tension from constant stretching and compression in the skin. The extent to which normal tissue movement requires MCs to withstand mechanical forces remains unclear. As cells experience tension, they can transmit mechanical forces from integrins to the nucleus through a contractile actomyosin (*82*). Perinuclear actin filaments connect to the nuclear envelope through the linker of nucleoskeleton and cytoskeleton (LINC) complex, allowing force transmission to deform the nucleus (*82, 83*). Previous work on non-immune cells in 2D culture systems demonstrated that nuclear flattening, dependent on perinuclear actin filament assembly and actomyosin contraction, activated nuclear transcription factors to induce G1-S transition (*84*). Our results indicate that the absence of Arp2/3-controlled actin filament branching causes shifts in the mechanical force balance within MCs. While Arp2/3-depleted MCs remain adherent, they lack lamellipodial protrusions and actin cortex integrity. Instead, MCs adopt a different mode of contractility-driven protrusiveness, forming lobopodia and cell blebs, most likely due to reduced actin cortex tension (*28, 37*). These substantial shifts in cytoskeletal forces could interfere with mechanical force transmission along the membrane-cortex interface, the cytoplasm, the perinuclear space, or a combination thereof, potentially contributing to the observed cell cycle arrest in G1/S phase and consequent cell death. Our data demonstrate that MC mechano-coupling to ECM requires the presence of Arp2/3-regulated branched actin filament networks. Therefore, we speculate that the Arp2/3 complex acts as a mechano-protective element, which safeguards MCs from mechanical forces, arising from cell movement or tissue mechanics. Thus, the Arp2/3 complex maintains the integrin-dependent MC positioning and cellular network organization in physiological tissues, which is required to mount functional immune responses.

In summary, our study emphasizes a significant physiological role of the Arp2/3 complex in preserving MC homeostasis *in vivo*, demonstrating that actin regulation is crucial for the survival and longevity of self-maintaining resident MCs. The maintenance of MC homeostasis necessitates not only continuous signaling through tissue-provided KIT ligand, but also Arp2/3-dependent actin regulation to resist integrin-mediated mechanical interactions with the tissue environment. This study provides insights into how actin filament branching impacts on immune cell function, particularly in the context of cell cycle regulation and tissue residency. Furthermore, our findings highlight that MC behavior is controlled by mechanical signals, paving the way for future investigations into the detailed underlying molecular or biophysical mechanisms, thus expanding the emerging field of mechanoimmunology (*85*).

## MATERIALS AND METHODS

### Study design

This study investigated the role of Arp2/3 complex-regulated actin filament branching for MC tissue residency. We generated conditional knockout mouse lines with MC-specific depletions for ARPC4, HEM1 and ARPC4 in combination with talin-1. Additionally, mice with constitutive deletion of WASP were examined. Mouse strains carried fluorescent reporters to image MC shapes *in vivo* or MC actin dynamics *in vitro.* Bone marrow (BM)-derived MCs (BMMCs) were generated from mice for *in vitro* experiments. To study endogenous MCs in tissues, fixed ear skins of differently aged mice were imaged with confocal fluorescence microscopy. Electron microscopy was utilized to characterize cell morphologies of *in vitro* cultured MCs. To mimic MC tissue dynamics, time-lapse video-based microscopy was established to observe MC dynamics and culture growth in 3D Matrigel or collagen I gels over 4 to 7 days. Retroviral fluorescence constructs were implemented for *in vitro* experiments to define cell cycle stages and determine the role of nuclear Arp2/3 in more detail.

### Mouse Models

Mouse breeding and husbandry were performed at the Max Planck Institute of Immunobiology and Epigenetics, Freiburg, in accordance with the guidelines provided by the Federation of European Laboratory Animal Science Association and as approved by German authorities (Regional Council of Freiburg). Mice were only used for organ removal after euthanasia by carbon dioxide exposure and thus not subject to experimental procedures and ethical approval according to §4 (3) Tierschutzgesetz. Mice were maintained in a conventional animal facility with a light-dark cycle of 14-10 hours at a temperature of 22°C ± 2°C and a relative humidity of 60% ± 5%. Standard food was available ad libitum for all animals. *Mcpt5-Cre* (*35*), *Arpc4^fl/fl^*(*33*), *Was^−/−^* (*86*), *Hem1^fl/fl^* (*43*), *Tln1^fl/fl^* (*87*)*, Tg(Ubow)* (*44*) and *Tg(Lifeact-GFP)* (*36*) mouse strains have been described elsewhere. *Mcpt5-Cre Arpc4^fl/fl^*, *Mcpt5-Cre Hem1^fl/fl^, Mcpt5-Cre Arpc4^fl/fl^Tln1^fl/fl^* mice and crosses with fluorescent reporter lines (*Tg(Ubow), Lifeact-GFP*) were on a C57BL/6J background. For all genotypes, age-and gender-matched female or male mice (aged 3 to 27 weeks) were used.

### Primary MC culture

To obtain MCs with connective tissue type characteristics, we cultured BMMCs according to a published protocol (*88, 89*). BM was isolated by flushing tibiae and femora with cold PBS. Isolated BM cells were maintained at 37°C and 5% CO_2_ in DMEM (4.5 g/L glucose, Gibco) supplemented with 10% heat-inactivated fetal calf serum (FCS), 10 U/ml penicillin, 10 µg/ml streptomycin, 2 mM L-glutamine, 25 mM HEPES, 1 mM sodium pyruvate, 1× non-essential amino acids (Gibco), 50 µM 2-mercaptoethanol and IL-3 (5% supernatant of murine IL-3-secreting WEHI-3 cells). To promote BMMC differentiation with connective tissue type characteristics, stem cell factor (SCF; 5% supernatant of murine SCF-secreting CHO transfectants) and IL-4 (1 ng/ml, PeproTech) were added to the medium (*89*). IL-4 and SCF supplementation to the culture medium enhanced Mcpt5-promoter activity and thus expression of CRE recombinase in BMMCs generated from Mcpt5-Cre mouse strains. BMMCs were used for experiments after 10 weeks of cultivation. Full differentiation into mature MCs was confirmed by expression of lineage-specific c-KIT (rat anti-CD117 FITC-conjugated, 1:200, Invitrogen) and FcεRI (armenian hamster anti-FcεRIα Alexa Fluor 647-conjugated, 1:200, Biolegend) measured by flow cytometry. BMMCs were passaged twice a week and kept at a concentration of 1 to 2.5 × 10^6^ cells/ml as a suspension culture. To assess the cell growth rate of BMMCs in media, 0.5 × 10^6^ viable cells were cultured as a suspension in untreated six well plates in 3 ml media supplemented with 10% IL-3 and 10% SCF after dead cell removal (Miltenyi-Biotec). Cells were counted at day 3, cultured in 10 ml fresh media with supplements and counted again at day 7.

#### IL-3, SCF and IgE production

IL-3 was produced by WEHI-3 cells (provided by R. Grosschedl, Max Planck Institute of Immunobiology and Epigenetics). SCF (KITLG) was produced by CHO transfectants (provided by G. Häcker, University of Freiburg). Anti-DNP IgE (clone SPE7) was produced by SPE-7 hybridoma NS1 cells (provided by M. Schmidt-Supprian, Technical University of Munich). WEHI-3 cells were cultured in DMEM (4.5 g/L glucose, 110 mg/L pyruvate; Gibco) supplemented with 10% heat-inactivated FCS, 10 U/ml penicillin, 10 µg/ml streptomycin, 2 mM L-glutamine and 50 µM 2-mercaptoethanol (*90*). Cells were kept at a concentration of 10^5^ cells/ml and medium was exchanged twice a week. For conditioned media production, cells were incubated for 3 to 4 days until the cell concentration reached 1 × 10^6^ cells/ml. Supernatant was collected, centrifuged for 10 min at 350 × g, filtered through a 0.45-μm filter (Nalgene Rapid-Flow) and kept at −20°C for long-term storage. CHO cells were kept at 37°C and 5% CO_2_ in OptiMEM^TM^ with Glutamax^TM^ (Life Technologies) supplemented with 10% heat-inactivated endotoxin-low FCS, 10 U/ml penicillin, 10 µg/ml streptomycin and 30 µM 2-mercaptoethanol. Cells were passaged twice a week and maintained as a confluent culture. For SCF production, cells were seeded at a concentration of 7 × 10^5^ cells per 15 cm cell culture plate (Greiner). When CHO cells reached 70% confluency, supernatant was collected and replaced by fresh medium. CHO supernatant harvest was repeated 3 to 4 consecutive days. The supernatant was centrifuged for 5 min at 500 × g, filtered through a 0.2-µm filter (Stericup plastic filter flasks, 0.2 μm) and kept at −20 °C.

### Human connective tissue MC and human MC lines

Human MCs were obtained with written consent of the patients and approved by the ethics committee of the Charité Universitätsmedizin Berlin (protocol code EA1/204/10, 9 March 2018). Cutaneous MCs were isolated from human foreskin tissue (*53*). Each MC culture originated from several (two to ten) donors. The experiments were conducted according to the Declaration of Helsinki principles. The skin was cut into strips and treated with dispase (24.5 mL per preparation, activity: 50 U/mL; Corning) at 4 °C overnight. The epidermis was removed, the dermis finely chopped and digested with 2.29 mg/mL collagenase (activity: 255 U/mg; Worthington), 0.75 mg/mL hyaluronidase (activity: 1000 U/mg; Sigma), and DNase I at 10 µg/mL (Applichem). The cell suspension was filtered stepwise (100 µm, 40 µm, 30 µm strainers). To purify the MCs, anti-human c-Kit microbeads and an Auto-MACS separation device were used (Miltenyi-Biotec), resulting in 98–100% MCs. Isolated skin MCs were measured by flow cytometry (double positive for c-KIT and FcεRI) and acidic toluidine blue staining (0.1 % in 0.5 N HCl). Skin MCs were cultured at 0.5 × 10^6^ cells/ml in basal Iscovés medium supplemented with 10% FCS, 1% P/S, SCF (100 ng/mL) and IL-4 (20 ng/mL), freshly provided twice weekly. MCs were used for experiments after 2.5 to 4.5 weeks. Two variant sublines of the human mast cell line HMC-1 were used (*91*). Human MC lines HMC-1.1 (Merck) and HMC-1.2 (Merck) were cultured in IMDM media (Sigma) with 1.2 mM alpha-thioglycerol (Merck), 10% heat inactivated fetal calf serum (Sigma) and 1× penicillin/streptomycin. The cells were kept at a concentration of 1-1.5 × 10^6^ cells/ml at 37°C and 5% CO_2_ and passaged twice a week.

### Flow cytometry analysis of MC maturation and cell death

To confirm MC maturation, flow cytometric analysis was performed. 1 × 10^6^ BMMCs per sample were used for staining and kept on ice. Antibodies and cells were diluted in PBS supplemented with 2% heat-inactivated FCS and 2 mM EDTA (FACS buffer). First, cells were blocked at 4°C for 15 min using anti-mouse CD16/CD32 antibody (1:250, BD Biosciences). Next, cells were stained for 30 min at 4°C with Alexa Fluor 647-conjugated anti-FcεRIα (1:200), FITC-conjugated anti-cKIT (1:200) and washed with FACS buffer. MCs were resuspended in FACS buffer and analyzed using a LSRIII^TM^ flow cytometer and FlowJo^TM^ software (BD Bioscience). Dead cells were excluded with DAPI staining. To monitor cell death during cultivation of WT and *Arpc4^−/−^* BMMCs, 10 to 14 weeks old BMMCs were cleared from dead cells with a dead cell removal kit (Miltenyi). Next, 0.5 × 10^6^ viable BMMCs were cultured and stained with Propidium iodide (1:100, Abcam) or DAPI (1:2000, Sigma) at day 0 and day 7 to assess the percentage of dead cells with flow cytometry.

### Determination of protein depletion efficiencies

To determine depletion efficiencies of Arpc4 and other Arp2/3 subunits in BMMC cultures, intracellular flow cytometry and western blot analysis were performed. For intracellular detection of ARPC2 protein expression by flow cytometry, 1 × 10^6^ BMMCs were fixed and permeabilized (Intracellular fixation kit, Cell Signaling Technology) according to the manufactures protocol. Unspecific binding sites were blocked as described above and then stained with anti-ARPC2 (1:50 in permeabilization buffer, Abcam) for 60 min at 4°C. Afterwards the cells were washed twice with permeabilization buffer and incubated with 488-conjugated anti-rabbit antibody (1:500, Invitrogen) for 60 min at 4°C, followed by two subsequent washing steps with permeabilization buffer. Cells were resuspended in FACS buffer for flow cytometric analysis.

For western blot analysis, 3 × 10^6^ BMMCs were lysed in a 1:3 ratio with Laemmli buffer (83 mM Tris pH 6.8, 13.3% glycerol, 1.3% SDS, 0.3 mg bromophenol blue, 0.47 M 2-mercaptoethanol) to ddH_2_O followed by 15–20 times of resuspension with a 23G needle (Braun). The samples were incubated at 95°C for 5 min and centrifuged 10 min at 13000 × g. Proteins were applied to a 12% gradient polyacrylamide gel, resolved by SDS-PAGE (BioRad) and transferred onto PVDF membranes (Millipore) via wet blot transfer. Nonspecific binding sites were blocked with block buffer (5% skim milk powder in Tris-buffered saline (TBS) containing 0.1% (v:v) Tween-20). Primary antibodies against the following proteins were incubated overnight in block buffer: ARPC4 (1:5000, Everest Biotech), ARPC2 (1:5000, Abcam), ARP3 (1:5000, Abcam), ARPC1A (1:5000, Abcam) and actin (1:5000, Santa Cruz Biotechnology). The membrane was washed three times in block buffer followed by a 1 h incubation with anti-goat/-rabbit HRP-conjugated secondary antibodies (1:5000, Agilent Dako) at RT. Proteins were visualized using Clarity Western ECL substrate (BioRad) and a ChemiDoc^TM^ Touch Gel Imaging System (Bio-Rad).

### Adhesion assay

To measure MC adhesion to different ECM components, an absorbance-based adhesion assay was performed. The method was performed as previously described with slight modifications (*12*). Microtiter plates (Greiner) were coated for 2 h at 37°C in triplicates with fibronectin (10 µg/ml, Sigma-Aldrich), Matrigel (50 µg/ml, Corning), laminin (5 µg/ml, Roche), bovine collagen I (30µg/ml, Advanced BioMatrix) or BSA as control (250 µg/ml) diluted in PBS. Plates were washed twice with PBS and blocked with 3% BSA in PBS for 30 min at 37°C. BMMCs were incubated for 1 h at 37°C in adhesion buffer (phenol red-free RPMI containing 10 mM HEPES, 0.25% BSA, and 2 mM CaCl_2_). Cells were washed once with PBS and seeded at 5 × 10^5^ cells/well in fresh adhesion buffer. Cells were allowed to settle for 10 min at 37°C, 5% CO_2_ and were then stimulated for 2 h with SCF (100 ng/ml). Next, plates were centrifuged upside down for 5 min at 60 × g to remove non-adherent cells. After washing once with pre-warmed PBS, plates were centrifuged again upside down. Adherent cells were fixed for 10 min with 4% PFA, followed by a washing step with PBS. Plates were centrifuged 2 min at 500 × g and adherent cells were stained by incubation with crystal violet (5 mg/ml in 2% ethanol) for 10 min at RT. Following two washing steps with tap water, plates were drained upside down and 1% SDS (in H_2_O) was added. Plates were put on an orbital shaker for 30 min to solubilize the crystal violet dye from lysed cells. Finally, the absorbance of the lysate was measured at 590 nm on a Synergy4 plate reader (Bio-Tek).

### Electron Microscopy

10 weeks old BMMCs were used for scanning electron microscopy. Dead WT and *Arpc4^−/−^* BMMCs were removed with a dead cell removal kit (Miltenyi-Biotec) and the viable cells were cultured in fresh BMMC culture medium with supplements overnight. BMMCs were adjusted to a concentration of 1 × 10^6^ cells/ml and fixed in suspension. Scanning electron microscopy was performed as described with slight modifications (*43*). BMMCs were fixed in suspension by adding formaldehyde (5% final concentration) and glutaraldehyde (2% final concentration) into growth medium. Afterwards suspensions were incubated on poly-l-lysine pretreated, round 12mm coverslips and washed twice with TE buffer (20 mM Tris, 1 mM EDTA, and pH 6.9). Dehydration with a graded series of acetone (10%, 30%, 50%, 70%, and 90%) was performed on ice for 10 min each, followed by two steps in 100% acetone at RT. Afterwards, samples were critically point dried (CPD 300, Leica, Wetzlar), fixed on aluminum stubs with plastic conductive carbon cement and covered with a gold palladium film by sputter coating (SCD 500 Bal-Tec, Liechtenstein). Examination was performed with a field emission scanning electron microscope Zeiss Merlin (Zeiss, Oberkochen) using both, the inlens SE (75%) and the Everhart Thornley HESE2 detector (25%) and an acceleration voltage of 5 kV.

### 3D Matrigel assay

To assess MC migration and longterm survival in 3D fibrillar matrices, BMMCs were embedded in Matrigel™ (Corning) for several days. Matrigel was used according to the manufacturer’s protocol. Dead BMMCs were removed (Miltenyi-Biotec) and viable cells were treated with 1 µg/ml anti-DNP IgE for 30 min at 37°C and 5% CO_2_. After washing with PBS, cells were suspended in BMMC culture medium with 5% WEHI-3 supernatant and 5% SCF-containing CHO supernatant at a concentration of 0.4 × 10^6^ cells/ml. Next, the BMMCs were mixed in a 1:1 ratio with Matrigel that was defrosted on ice for 24 h prior use. The mixture was gently resuspended for 60 sec and 50 µl were added in a 96 well image lock plate (Sartorius) on ice. The 96 well plate was centrifuged at 75 × g, 4°C for 3 min to enable a homogenous cell distribution and then incubated at 37°C and 5% CO_2_ for 30 min for Matrigel polymerization. Afterwards, 250 µl BMMC culture medium with 2 × concentrated growth factors (10% IL-3-containing WEHI-3 and 10% SCF-containing CHO supernatant) was added on top of the gel. Live cell imaging was performed using an Incucyte S3 live-cell analysis system (Sartorius). Each well was imaged in a 15 min interval for up to 7 days with the image lock module and a 10× or 20× objective. Red and green fluorescent signals were acquired with the in-built dual color module 4614. For cell death analysis sytox orange (1:1000, Thermo Fisher) was added to the medium on top of the polymerized gel. When required, anti-DNP IgE sensitized BMMCs were pre-incubated with inhibitors or blocking antibody: 50 µM Y-27632 (Merck), 50 µM blebbistatin (Merck), 200 µM CK666 (Merck) and 5 µg/ml anti-ITGB1 (BD Bioscience) in BMMC culture media at 37°C and 5% CO_2_ for 30 min. As controls, cells were treated with DMSO for blebbistatin, H_2_O for Y-27632 or with the inactive CK666 variant CK689 (Merck). After chemical incubation, the cells were washed twice with PBS, resuspended in BMMC culture medium and embedded in Matrigel. The chemicals were also added in the culture medium on top of the gel to enable a stable concentration during the live cell imaging experiment. For cell starvation experiments, anti-DNP IgE sensitized BMMCs were washed with 1× PBS, resuspended in PBS followed by embedding in the Matrigel. After Matrigel polymerization, 250 µl PBS with sytox orange were added on top of the solidified gel. To obtain a higher magnification of cells in the Matrigel, live cell imaging with a confocal spinning-disc microscope (Zeiss) was performed and the Matrigel preparation procedure adapted. Anti-DNP IgE sensitized BMMCs were adjusted to 1 × 10^6^ cells/ml and if required treated with chemicals as described above. After washing, 25 µl of the cell suspension was mixed with 25 µl Matrigel on ice. 10 µl were loaded on an µ-angiogenesis slide (Ibidi) and incubated for 10 min at 37°C and 5% CO_2_ for gel polymerization. 45 µl BMMC media were added on top and the slide was kept in the incubator at 37°C and 5% CO_2_ overnight before imaging. For nucleus visualization, Hoechst (1:5000, Merck) was added to the BMMC medium on top of the gel and incubated at 37°C and 5% CO_2_ for 3 h before imaging. For imaging an EC Plan-Neofluar 40×/1.30 Oil (Zeiss) or a Plan-Apochromat 63×/1.40 Oil (Zeiss) objective was used and the cells were kept in a stage top incubator (TokaiHit) at 37°C and 5% CO_2_ to enable physiological conditions. In addition, a z-stack of 1 µm steps was set to ensure good imaging quality without causing phototoxicity. Live cell imaging intervals were between 1–3 min for 2–45 min imaging time in total.

### 3D collagen I gel assay

To monitor MC dynamics and long-term survival in 3D collagen gels, BMMCs were embedded in bovine collagen I gels (Advanced BioMatrix). Cells were prepared and imaged as described above for 3D Matrigel. After dead cell removal and anti-DNP IgE incubation, cells were adjusted to 0.8 × 10^6^ cells/ml or 1.2 × 10^6^ cells/ml (20× objective) in supplemented BMMC culture media. To prepare the collagen I gel, pre-chilled NaHCO_3_, pre-chilled 10× MEM (minimal essential medium, Sigma) and pre-chilled bovine collagen I (Advanced BioMatrix) were mixed together with a final pH between 7 and 8. The collagen suspension was mixed carefully with the cells in a ratio of 4:1 and 50 µl were added in a 96 well image lock plate (Sartorius) on ice. The centrifuged 96 well plate was incubated for 1 h at 37°C and 5% CO_2_ to solidify the collagen gel. The cells were imaged with an Incucyte S3 live-cell imaging system as described above. For confocal imaging, the cell concentration was adapted to 1 × 10^6^ cells/ml and 10 µl of the cell suspension:collagen mix was added to an µ-angiogenesis slide.

### Ear skin whole mount immunofluorescence analysis

Ears were cut at the base and subsequently split into ventral and dorsal halves. Ventral ear sheets were fixed in 1% PFA overnight at 4°C on a shaker. Ear explants of Ubow mice were fixed for only 4 h in 1% PFA at 4°C to reduce quenching of the weak CFP fluorescence signal. Ear slices were permeabilized with block buffer (0.25% (v/v) Triton^TM^ X-100 in PBS with 1% BSA) and incubated for 1 h at 4°C while gently shaking. Next, immunofluorescence staining was performed at 4°C on a rocker at all steps. Between each step the samples were washed three times for 15 min in block buffer. Primary antibodies or directly fluorescent reagents were applied overnight at 4°C. If required, this was followed by a 4 to 6 h incubation at 4°C with fluorescently labeled secondary antibodies. Tissue samples were mounted onto glass slides and covered by glass cover slips (the dermal side facing the cover slip) with Fluoromount-G (SouthernBiotech). The following antibodies were used: anti-ACTA2 Cy3-conjugated (1:500, Sigma-Aldrich), anti-collagen IV (1:500, Abcam), Anti-rabbit Alexa Fluor 405-conjugated. Avidin was used to visualize dermal MCs (1:10000, FITC-conjugated) To receive a more detailed morphology of cellular structures, the endogenous YFP and CFP signals of Ubow mice strains were amplified with Dylight™ 488-conjugated anti-GFP antibody (1:1000, Rockland). *Z*-stacks were acquired (indicated in the respective analysis section and/or figure legend). Standard ear whole mount microscopy was performed using a LSM 780 microscope (Zeiss). 2 × 2 tile images were acquired covering 30–44 µm thick *z*-stacks with a step size of 1.5–2 µm using a Plan Apochromat 20×/ 0.8 air objective (Zeiss) with 0.6 scan zoom. Tiled images were stitched with 10% overlap during post-processing with ZEN blue software (Zeiss). For detailed morphology images of Ubow mice, a scan zoom of 1 was used with an Plan Apochromat 20×/0.8 air objective (Zeiss) or a Plan Apochromat 40×/1.4 Oil DIC M27 objective (Zeiss).

### Adhesion cell death assay

Tissue culture treated 24 well plates were coated with fibronectin (10 µg/ml, Sigma) and incubated for 30 min at 37°C and 5% CO_2_. Each well was washed twice with PBS and filled with 150 µl BMMC culture media, supplemented with 10% IL-3 and 10% SCF. Anti-DNP IgE sensitized BMMCs were adjusted to a cell concentration of 1 × 10^6^ cells/ml in BMMC culture medium with supplements. 1 × 10^5^ cells were added into each well and incubated at 37°C and 5% CO_2_. Cell death measurements were performed after 24 h and 48 h. Therefore, cells were transferred in 1.5 ml tubes and adherent cells were detached with EDTA (2 mM in PBS) for 30 sec at 37°C. The samples were centrifuged for 5 min at 240 × g. The pellets were resuspended in BMMC culture medium with propidium iodide (PI) (1:100, Abcam) and the cell death was quantified by counting PI positive cells using flow cytometry.

### Purification of cellular fragments, mass spectrometry, LC-MS and MS data analysis

Details are provided in the Supplementary Materials.

### Retrovirus production and BMMC transduction

For cell cycle analysis, the pBMN-FUCCI plasmid was generated by replacing the lentiviral backbone pBOB from the original plasmid with the retroviral pBMN-Z backbone. The original plasmid pBOB-EF1-FastFUCCI-Puro was a gift from Kevin Brindle & Duncan Jodrell (Addgene plasmid #86849; RRID: Addgene_86849) (*92*). For nuclear Arp2/3 inhibition, the pBMN-mScarlett-Arpin-NLS plasmid was provided by R. Grosse. For retrovirus production, 2.5 × 10^6^ PlatE cells were plated on 10 cm tissue culture dishes in DMEM culture medium. The following day PlatE cells were transfected with pBMN-plasmids using a Lipofectamin 3000 kit (Invitrogen) according to the manufactures protocol. Retroviral supernatant was collected every 24 h for up to 4 days and stored at 4°C. For retroviral transduction, 9 to 12 days old BMMCs were adjusted to a cell concentration of 0.8 × 10^6^ cells/ml, centrifuged and resuspended in retroviral supernatant with 5% SCF and 10 µg/ml Polybrene. The cells were plated on a 6 well cell culture plate and centrifuged at 1500 × g and 35°C for 3 h. After centrifugation the cells were incubated at 37°C and 5% CO_2_ for 2 h, followed by an overnight incubation in BMMC culture medium with 5 % IL-3, 5 % SCF and 10 % viral supernatant. The next day, the cells were washed and resuspended in BMMC media with 5 % IL-3 and 5 % SCF and cultured at 37°C and 5% CO_2_ until the transduced cells were sorted with a MoFlow sorter (Beckmann Coulter) by reporter expression. The sorted transduced BMMCs were further cultured and used for experiments at 10 weeks of age.

### Analysis of MC morphologies and migration in 3D Matrigel

To assess the morphology of WT and *Arpc4^−/−^* BMMCs in 3D Matrigel, *n=*50 cells of each independent experiment were randomly selected and the area was manually measured with ImageJ (V2.1.0/1.53c) (*93*). Circularity values per cell were analysed with the shape descriptor “circularity” = 4π × (area/perimeter^2^). Live cell tracking of MCs in 3D Matrigel was performed using the manual tracking option of Imaris 9.1.2 (Bitplane). Cells were randomly chosen and tracked manually through all time frames. The average cell speed was calculated by dividing the total track length by the total time of imaging in minutes.

### Calculation of MC density in tissues

To determine the number of MCs in the mouse ear skin tissue, ear skin whole mounts of WT (*n*=3) and *Arpc4^−/−^* (*n*=3) mice in age groups of 3 weeks, 11–13 weeks and 27 weeks were quantified. 2 × 2 tile images were acquired as described above. Three imaging fields of view, displayed as maximum intensity projection were analysed per mouse and the cell number was quantified with ImageJ (V2.1.0/1.53c). Therefore, the avidin signal (MCs) was masked using the plugin morpholibJ (*94*). The masked cells were automatically counted. Clustered cells were manually counted and degranulated MCs were manually removed from the analysis. The MC density was displayed per analyzed tissue area in mm^2^.

### Cellular fragment quantification

The number of fragments in 3D Matrigel and 3D collagen gel were quantified with ImageJ (V2.1.0/1.53c) and Imaris 9.5.1. (Bitplane). Therefore, 3 imaging fields of view of each independent experiment were quantified. A defined volume of 1.41 × 10^6^ µm^3^ was generated for all imaging fields of view with Imaris, the number of fragments were quantified with ImageJ and displayed per analyzed volume in mm^3^. To determine the number of cellular fragments in the ear skin tissue, ear skin whole mounts of Ubow WT (*n*=4) and Ubow *Arpc4^−/−^* (*n*=4) mice in age groups of 18-21 weeks were quantified with Imaris. Images were acquired with a Plan Apochromat 20×/ 0.8 air objective (Zeiss) with scan zoom 1. Three imaging fields of view, displayed as maximum intensity projection were analyzed per mouse and the number of avidin positive and GFP (Ubow) positive fragments were quantified with the spot function. The number of fragments was displayed per analyzed tissue area in mm^2^.

### Determination of MC numbers, cell cycle stages and cell death in 3D Matrigel

To quantify the cell number, cell death rate and cell cycle stages of BMMCs in Matrigel and collagen, snapshots of different time points were analyzed with ImageJ (V2.1.0/1.53c). For the cell number quantification, the cells were both manually as well as automatically counted. Automatically, the cells were masked with the plugin morpholibJ, cell clusters were automatically separated by a watershed function and cells were counted. Automatic cell counting with cells expressing the arpin construct was performed only with the red fluorophore channel to exclude non-transduced cells from the analysis. To evaluate the number of dead cells, the phase contrast channel and the fluorophore channel displaying the cell death were merged and the dead cells were manually counted. To analyze the numbers of cell cycle stages in 3D, the phase contrast channel was merged with the fluorophore channels displaying the different cell cycle stages (G1/S in red, G2M in green). The numbers of cells in different cell cycle stages could be distinguished by colour and quantified manually.

### Cell division and single cell fate mapping analysis in 3D

The percentage of dividing cells in 3D was analysed with ImageJ (V2.1.0/1.53c). First, the total cell number per independent experiment was counted manually. Second, the number of cell divisions was evaluated manually by green nucleus colour and quantified as percent from the total cell number. For the cell fate mapping analysis in 3D Matrigel, 35 cells were randomly selected and manually followed over 96 h. Every 3 h the cell cycle stage was evaluated and displayed as a heatmap.

### Statistical analysis

Analyses were performed using Prism software (GraphPad Software, Inc.Version 9.3.1). If not indicated otherwise, comparisons for two groups were evaluated using a two-tailed unpaired student’s *t* tests after confirming that samples fulfil the criteria of normality. The D’Agostino & Pearson normality test was performed for group sizes over 10, for group sizes under 10 the Shapiro-Wilk normality test was performed. For *in vitro* experiments biological replicates were cell cultures generated from different mice and independent experiments were performed at different time points. For not-normally distributed data, non-parametric Mann-Whitney U tests were used. Stars indicate significance (\**P* ≤ 0.05, \*\**P* ≤ 0.01, \*\*\**P* ≤ 0.001). NS indicates non-significant difference (*P* > 0.05). For further statistical details, see Supplementary Table S2.

## Supplementary Materials

Materials and Methods Figs. S1 to S6

Tables S1 to S3

Legends to Movies S1 to S6 References

## Acknowledgments

We thank R. Wedlich-Söldner, K.A. Siminovitch, A. Roers, S. Monkley and D. Critchley for kindly providing mice for this study, K. Lucht for help with molecular biology work, members of the MPI Imaging Facility for assistance with imaging, the flow cytometry facility for cell sorting, I. Brentrop for EM sample preparation, and members of the Lämmermann lab for helpful discussions; **Funding:** This work was supported by the Max Planck Society (T.L.), IZKF, Medical Faculty, University of Münster (T.L.), the Deutsche Forschungsgemeinschaft DFG (GR 2111/13-1, R.G.; BA-3769/3, BA-3769/4, M.B.); the European Union—NextGenerationEU through the Italian Ministry of University and Research under PNRR—M4C2-I1.3 Project PE_00000019 “HEAL ITALIA”, CUP H43C22000830006 of University of Milano Bicocca (M.I.). The views and opinions expressed are those of the author only and do not necessarily reflect those of the European Union or the European Commission. Neither the European Union nor the European Commission can be held responsible for them. **Author contributions:** LK: Conceptualization, Investigation, Methodology, Validation, Formal Analysis, Visualization, Writing – Original Draft, TEBS, MI, MB, SU, RG: Resources, Methodology, MMi, MMü, GM: Investigation, KMG: Visualization, TL: Conceptualization, Funding Acquisition, Project Administration, Supervision, Investigation, Visualization, Writing – original draft, all authors: Writing – review and editing; **Competing interests:** Authors declare no competing interests.; and **Data and materials availability:** All data is available in the main text or the supplementary materials. *Mcpt5^CRE^*mice are available through Axel Roers (Universitätsklinikum Heidelberg), *Was^-/-^*mice are available through Katherine A. Siminovitch (University of Toronto), *Hem^fl/fl^* mice are available through Theresia E.B. Stradal (Helmholtz Center for Infection Biology, Braunschweig).

