## Supplementary Figures 1-6, Legends to Supplementary Videos 1-6 for "Arp2/3 complex-dependent actin regulation protects the survival of tissue-resident mast cells"

Lukas Kaltenbach *et al.*

**Supplementary Information:**

Materials and Methods
Figs. S1 to S6
Tables S1 to S3
Legends to Movies S1 to S6
References

### Materials and Methods

#### *Purification of cellular fragments and mass spectrometry*

The cellular fragments were purified by multiple centrifugation steps and subsequently analyzed with mass spectrometry. Treated 24-well plates were coated with fibronectin (10 µg/ml in PBS, Sigma Aldrich) in PBS for 30 min at 37°C and 5% $CO_2$ and afterwards washed twice. WT and *Arpc4*$^{-/-}$ BMMCs were cleared from dead cells using a dead cell removal kit (Miltenyi Biotec), incubated with anti-DNP IgE as described above and adjusted to a final concentration of 0.2 × $10^6$ cells/ml in BMMC media with 10% IL-3 and 10% SCF. 1 × $10^5$ BMMCs were added per well in a total volume of 500 µl BMMC culture medium with 10% IL-3 and 10% SCF followed by 48 h incubation at 37°C and 5% $CO_2$. For the fragment purification process, the supernatant of the wells was taken up carefully and transferred into 1.5 ml tubes. The tubes were centrifuged at 300 × g for 10 min at RT and the supernatant (S1) was transferred into a fresh 1.5 ml tube. S1 was then centrifuged at 5000 × g for 10 min and the supernatant (S2) was transferred in a new 1.5 ml tube. S2 was again centrifuged at 5000 × g for 10 min and the pellet was resuspended in Tyrode's buffer (135 mM NaCl, 5 mM KCL, 5.6 mM glucose, 1.8 mM $CaCl_2$, 1 mM $MgCl_2$, 20 mM HEPES-NaOH, pH to 7.3) and filtered with a 5 µm filter (Sysmex) in protein low-bind tubes (Eppendorf). As a control for successful fragment purification the pellet was measured by flow cytometry for FSC and SSC. To visualize the purified fragments, Lifeact-GFP expressing BMMCs were used. The pellet was directly embedded in Matrigel and the fragments could be visualized with a confocal spinning-disk microscope (Zeiss). Therefore, a Plan-Apochromat 100×/1.40 Oil DIC M27 objective (Zeiss) was used and a *z*-stack with a step size of 1 µm. For mass spectrometry the particles were dissolved in 55 µl EP lysis buffer (100 mM Tris-HCl, pH 7.6, 1% SDS, 1.5 mM $MgCl_2$, 0.2 mM EDTA, 10 mM TCEP, 40 mM chloroacetamide) and treated as follows. After an initial incubation on ice (10 min) samples were heated at 90°C (5 min) and cooled on ice. This was followed by adding 0.5 µl Benzonase (100 U/µl, EMD Millipore) and subsequent sonication (Diagenode Bioruptor Plus, five cycles, 30 s "on", 30 s "off") and a final incubation at 25°C (10 min). Next, samples were subjected to paramagnetic bead-based single-pot, solid-phase-enhanced sample-preparation (SP3). 7.5 µl Sera-Mag Carboxylate-modified beads (Cytiva) suspension (20 µg/µl stock in LC-MS $H_2O$; 1:1 mixture of carboxylate-modified Sera-Mag Speed Beads A (hydrophilic) and B (hydrophobic), respectively) was added to the samples and protein binding was induced by addition of neat acetonitrile to a final concentration of 73% followed by 15 min incubation at RT (800 rpm, Eppendorf mix mate). Beads were collected by incubation for 15 min at RT on an in-house constructed magnetic rack. Washing steps consisted of three washes with 400 µl 70% ethanol and one wash with 400 µl neat acetonitrile. Finally, beads were transferred to 0.5 ml low-bind tubes and briefly air-dried before reconstitution in 50 µl 50 mM ammonium bicarbonate (Honeywell) containing 0.02% Rapigest (Waters). Proteolytic digestion was started with 100 ng LysC (Wako) and incubation at 37°C for 2 h (Thermomixer, 1400 rpm). Subsequently, 500 ng sequencing grade trypsin (Promega) were admixed followed by incubation at 37°C for 12 h (Thermomixer, 1400 rpm). Beads and tryptic peptides were efficiently resuspended by sonication (Bandelin Sonorex water bath sonicator, 1 min) and concentrated *in vacuo* to a final volume of 5 µl. Then samples were vortexed briefly and subjected to sonication as described above. Rebinding of peptides to magnetic beads was induced by addition of neat acetonitrile to a final concentration of 90% followed by 15 min incubation at RT (800 rpm, Eppendorf mix mate). Beads were collected by incubation for 20 min at RT on an in-house constructed magnetic rack, washed once with 200 µl neat acetonitrile and briefly air-dried. Then, we conducted a two-step elution of tryptic peptides from the beads. They were supplied with 40 µl LC-MS $H_2O$ (Pierce), resuspended by sonication (Bandelin Sonorex water bath sonicator, 2 min) and collected on a magnetic rack (15 min, RT). The supernatant was transferred to a fresh 0.5 ml low binding tube. The beads were further extracted with 20 µl 0.1 % trifluoroacetic acid (Bandelin Sonorex water bath sonicator, 2 min) and collected (magnetic rack, 15 min, RT). Extracted peptides were combined (60 µl volume), transferred to 96 well plates and reduced *in vacuo* to a volume of ~2 µl followed by addition of 1.5 µl 5% acetonitrile in 0.1% formic acid and 6 µl of 0.1% formic acid prior to nanoLC-MS analysis.

**LC-MS analysis**

An Orbitrap Exploris™ 480 coupled to an EASY-nLC 1200 nUHPLC (both Thermo Fisher Scientific) was used for all measurements. General LC-MS setup was essentially as described (*1*). Samples were injected three times (once with a 30 min, once with a 120 min and once with a 180 min nLC-MS method). The gradient for the 30 min nLC-MS method was: start (5%), 2 min: 10%, 18 min: 60%, 1 min: 90% B (150 nl/min flow rate). This was followed by a "wash out step": 4 min: 90% B buffer and a 1.5 min inverse gradient from 90% to 0% B buffer (flow rates 500 nl/min). The gradient for the 120 min nLC-MS method was: start (0%), 5 min: 4%, 90 min: 30%, 10 min: 40%, 2 min: 80% B (300 nl/min flow rate). This was followed by a "wash out step": 5 min: 80% B buffer and a 2 min inverse gradient from 80% to 2% B buffer (flow rates 500 nl/min). The gradient for the 180 min nLC-MS method was: start (2%), 5 min: 5%, 150 min: 30%, 12 min: 40%, 3 min: 80% B (300 nl/min flow rate). This was followed by a "wash out step": 5 min: 80% B buffer and a 2 min inverse gradient from 80% to 2% B buffer (flow rates 500 nl/min). All measurements were carried out in data-dependent mode with full MS resolution of 120,000 (30 min method) or 60,000 (120 min and 180 min method) at m/z 200, AGC target 300%, mass range 350-1650 m/z, IT of 45 ms (120 min method)/60 ms (30 min and 180 min method) and dynamic exclusion of 15 s (30 min method)/30 s (120 min method)/36 s (180 min method). MS/MS spectra acquisition was done with a resolution of 30,000 (30 min method)/7,500 (120 min and 180 min method), AGC target 200%, and 1.5 m/z isolation windows at a normalized collision energy of 30 together with and IT of 50 ms (30 min method)/22 ms (120 min method)/30 ms (180 min method). General MS conditions were 2.1 kV positive ion spray voltage, 275°C ion transfer tube temperature, 1% sweep gas and RF lense voltage set to 45%.

**MS data analysis**

MaxQuant (v. 1.6.14.0) employing standard parameters (*1*) with exemptions detailed below was used to identify peptides and final proteins identification role-up (both at 1%FDR). Technical replicates (LC-MS injections) were collapsed into one experimental group (for each biological replicate). MS raw data (.RAW files) were searched simultaneously with the target-decoy standard settings against the Uniprot *Mus musculus* database (Uniprot_reviewed+Trembl including canonical isoforms; downloaded on August 5th, 2020) appended with an in-house curated FASTA file containing commonly observed contaminant proteins based on MaxQuant's contaminants list (*2*). Initial mass tolerance was 20 ppm followed by 4.5 ppm for main search and fragment tolerance of 25 ppm. Trypsin/P was used as enzyme and up to 2 missed cleavages and a minimal peptide length of six were allowed. Carbamidomethylation of cysteine was set as fixed modification. Variable modifications included oxidation (M), deamidation (N, Q) and acetylation of protein N-termini. Inter-sample relative abundance was determined using MaxLFQ (*3*)with disabling the match between runs option (minimal LFQ ratio count at 1). Intra sample abundance was approximated by iBAQ score. Peptide and protein FDR were kept at 1 %. All other settings were kept at default. The proteingroups.txt output file was employed for further analysis. Briefly, contaminant proteins, reverse (decoy) data base hits, and proteins only identified by site entries were first filtered out. For the downstream analysis, protein hits were sorted by LFQ Intensity in descending order. The top 50 hits were analyzed for functional annotation with DAVID ontology (*4, 5*).

**Fig. S1.**

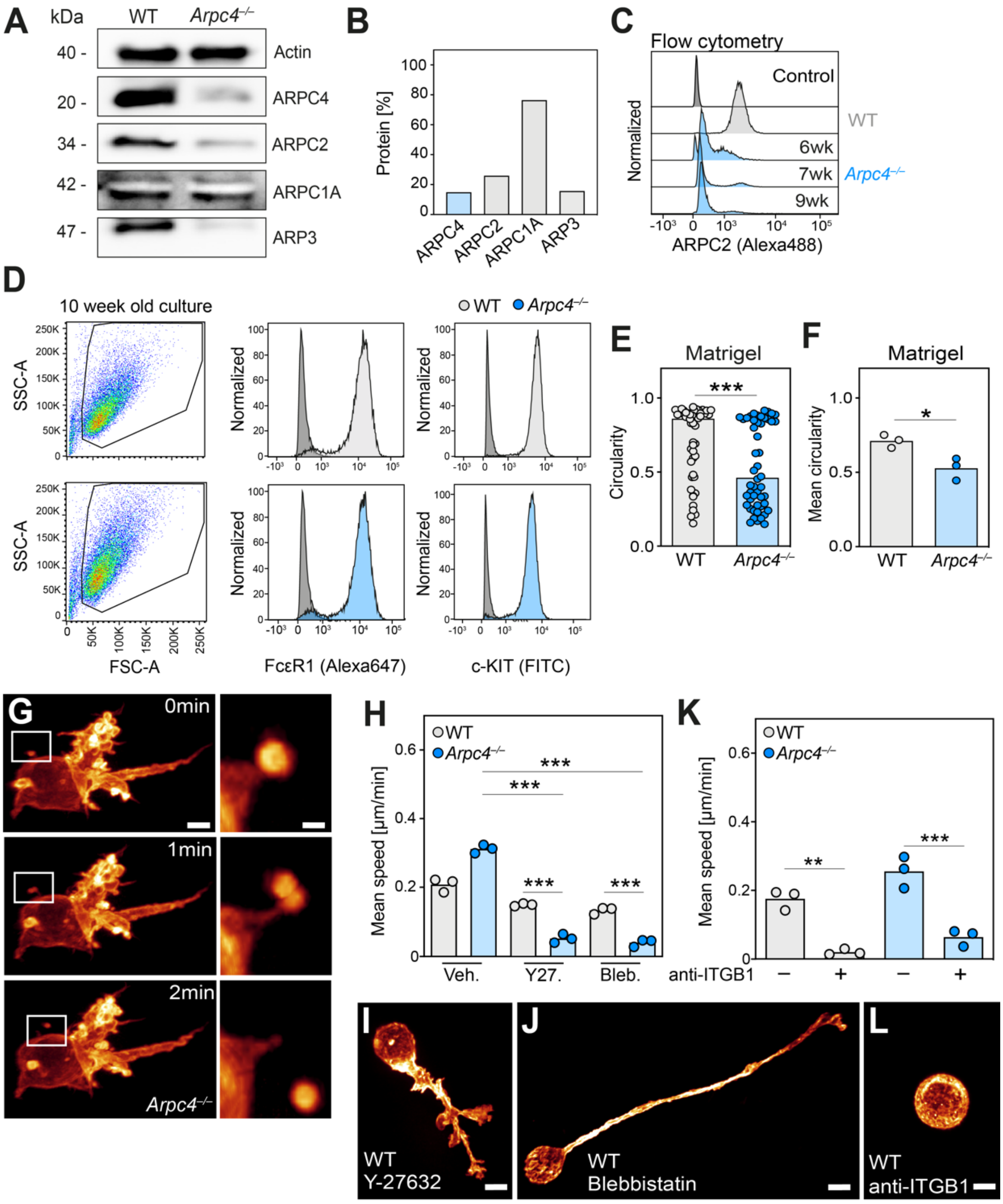


**Fig. S1: Characterization of *Arpc4* knockout efficiency, MC maturation and migration.**

**(A–C)** *Arpc4*$^{-/-}$ BMMCs were generated from *Arpc4*$^{\Delta MC}$ mice (Fig. 1A). Efficiency of ARPC4 depletion and downregulation of other ARP2/3 subunits was confirmed by immunoblot analysis of cell lysates (A, B) and intracellular flow cytometry using anti-ARPC2 antibody (C). Actin was used as loading control (A) and the percentage of protein expression was quantified by normalization to the loading control (B). Intracellular flow cytometry of BMMCs, gated by FSC/SSC (see also D), confirmed the result from the immunoblot of ARPC2. Secondary antibody control (dark grey) is shown on top (C). **(D)** Comparable expression of MC maturation markers c-KIT and FcεR1 in WT and *Arpc4*$^{-/-}$ BMMCs. Gating for BMMCs by FSC/SSC. **(E, F)** Quantification of cell roundness (circularity) was performed for WT and *Arpc4*$^{-/-}$ BMMCs in 3D Matrigel after t=36 h. (E) *N*=50 cells per genotype were quantified per independent experiment; bars display the median; ****P*≤0.001, *U* test. (F) Mean values of *n*=3 independent experiments from 3 biological replicates; bars display the mean; **P*≤0.05, *t* test. **(G)** Spinning-disk confocal microscopy of cellular fragment loss from the cell body of a Lifeact-GFP expressing *Arpc4*$^{-/-}$ BMMC in 3D Matrigel. A time-sequence from live-cell imaging over 2 min is displayed. **(H–J)** To address the role of with actomyosin contraction, WT and *Arpc4*$^{-/-}$ BMMCs were treated with Y-27632 (Y27.) or blebbistatin (Bleb.). (H) Quantification of the mean cell speed per independent experiment (*n*=3 independent experiments from 2 biological replicates) from Fig. 1K. (I, J) Representative morphologies of Lifeact-GFP (glow) expressing WT BMMCs in 3D Matrigel upon inhibitor treatment. **(K, L)** To interfere with integrin β1 (ITGB1)-mediated adhesion, WT and *Arpc4*$^{-/-}$ BMMCs were treated with anti-ITGB1 antibody. (K) Quantification of the mean cell speed per independent experiment (*n*=3 independent experiments from 2 biological replicates) from Fig. 1M. (L) Representative morphology of Lifeact-GFP (glow) expressing WT BMMCs in 3D Matrigel upon anti-ITGB1 treatment. Scale bars: 5 µm, 1 µm (zoom-in) (G), 15 µm (I, J, L). (H, K) Bars display the mean; ***P*≤0.01, ****P*≤0.001, Tukey´s multiple comparison (posthoc one-way Anova test).

**Fig. S2.**

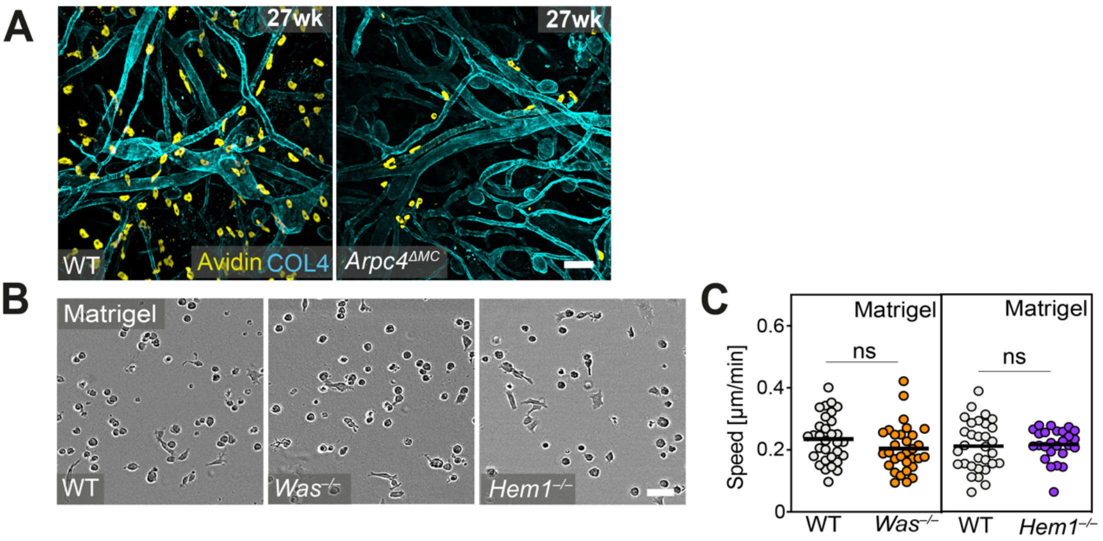


**Fig. S2: Depletion of WASP and Hem1 do not affect MC migration dynamics in 3D Matrigel.**

**(A)** Ear skin whole mount tissues of adult *Arpc4*$^{\Delta MC}$ mice and littermate controls. Dermal MCs were immuno-stained with fluorescent avidin (yellow) in relation to collagen IV (COL4)-expressing basement membrane structures (blue), which visualize all blood and lymph vessels, nerve bundles and fat cells. **(B, C)** BMMC cultures from *Was*$^{-/-}$ and *Hem1*$^{\Delta MC}$ mice were generated and studied in 3D gels. (B) Representative phase-contrast brightfield images of WT, *Was*$^{-/-}$ and *Hem1*$^{-/-}$ BMMC moprhologies in 3D Matrigel. (C) Quantification of the cell speed over 36 h is shown. Dots in the graph are values of individual cells ($N$=30 randomly chosen cells per genotype). Bars display the mean; ns, non-significant, *t* test. Scale bars: 70 µm (A), 50 µm (B).

Fig. S3.

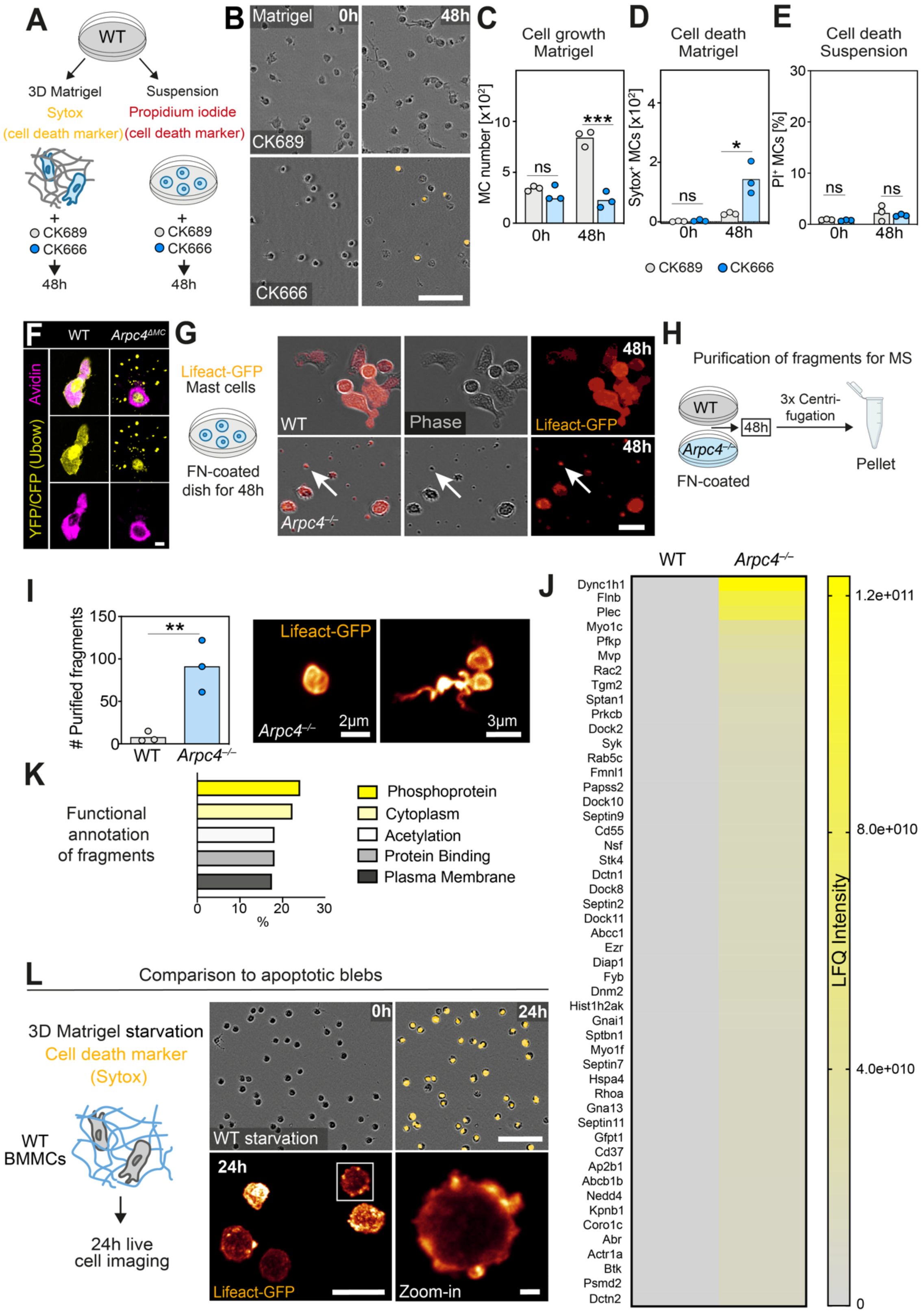


**Fig. S3: Effects of pharmacological Arp2/3 inhibition and characterization of *Arpc4*$^{-/-}$ MC fragments.**

**(A)** WT BMMCs were treated with the Arp2/3 inhibitor CK666 or its inactive form CK689 and cell death measured in 3D Matrigel or in suspension. Cells undergoing cell death were detected with Sytox (orange) in 3D Matrigel or with propidium iodide (PI) in suspension. **(B-D)** Assessment of cell death in

3D Matrigel. (B) Brightfield images of CK666- and CK689-treated WT BMMCs in the presence the cell death marker Sytox (orange) ($t$=0 h, $t$=48 h). (C, D) Quantification of MC proliferation and cell death at time points 0 h and 48 h. Each dot represents one independent experiment per treatment ($n$=3 independent experiments from 2 biological replicates). (C) Bars display the median (0 h) or mean (48 h). ns, non-significant, $U$ test (0 h). ***$P \leq 0.001$, $t$ test (48 h). (D) Bars display the mean; *$P \leq 0.05$, $t$ test (48 h). **(E)** Quantification of cell death in suspension by measuring the percentage of PI-positive cells with flow cytometry. $n$=3 independent experiments per genotype. Bars display the mean; ns, non-significant, $t$ test. **(F)** Representative images of dermal MCs in the ear dermis of WT Ubow or $Arpc4^{\Delta MC}$ Ubow mice (Fig. 3I). Ear skin whole mounts were stained with fluorescent avidin (purple) and anti-GFP to amplify YFP/CFP signals of the Ubow reporter (yellow). Amplified Ubow signals display cell fragments in tissue areas around *Arpc4*-deficient MCs. **(G, H)** Actin-containing fragments of $Arpc4^{-/-}$ BMMCs adhered to fibronectin (FN)-coated 2D dishes over 48 h, were then purified and prepared for mass spectrometry (MS) analysis. (G) Representative images of Lifeact-GFP expressing WT and $Arpc4^{-/-}$ BMMCs. $Arpc4^{-/-}$ MC adhesion to FN-coated dishes results in fragmentation formation (white arrow) over 48 h. (H) Scheme of the cell fragment purification process. **(I)** Fragment quantification of WT and $Arpc4^{-/-}$ BMMCs by flow cytometry. Each dot represents the total number of purified fragments of one independent experiment ($n$=3 independent experiments from 2 biological replicates). Bars display the mean; **$P \leq 0.01$, $t$ test. Spinning-disk confocal imaging of representative purified Lifeact-GFP positive cell fragments from $Arpc4^{-/-}$ MCs. Fragments were embedded in 3D gels. **(J, K)** Purified fragments of WT and $Arpc4^{-/-}$ BMMCs were analyzed with MS. (J) Heatmap of the top 50 protein hits sorted by LFQ intensity. As WT BMMCs formed hardly any fragments, WT samples were below detection limit. (K) Functional annotation analysis of top 50 protein hits of $Arpc4^{-/-}$ BMMC fragments performed with DAVID ontology. **(L)** Comparison of cell fragments to apoptotic cell blebs. WT BMMCs in 3D Matrigel were starved to induce apoptotic cell death. WT BMMCs were starved with PBS and imaged for 24 h in 3D gels. Representative phase-contrast brightfield microcopy images with Sytox cell death marker (orange) (t=0 h, t=24 h). Spinning-disk confocal imaging of starved Lifeact-GFP WT BMMCs at t=24 h in 3D Matrigel. WT BMMC in the zoom-in displays bleb-like structures connected with the plasma membrane. Apoptotic cell induction and blebbing did not lead to the visible release of cell fragments as observed for $Arpc4^{-/-}$ MCs. Scale bars: 80 µm (B), 5 µm (F), 50 µm (G), phase-contrast: 80 µm, Lifeact: 20 µm, 2 µm (zoom-in) (L).

**Fig. S4.**

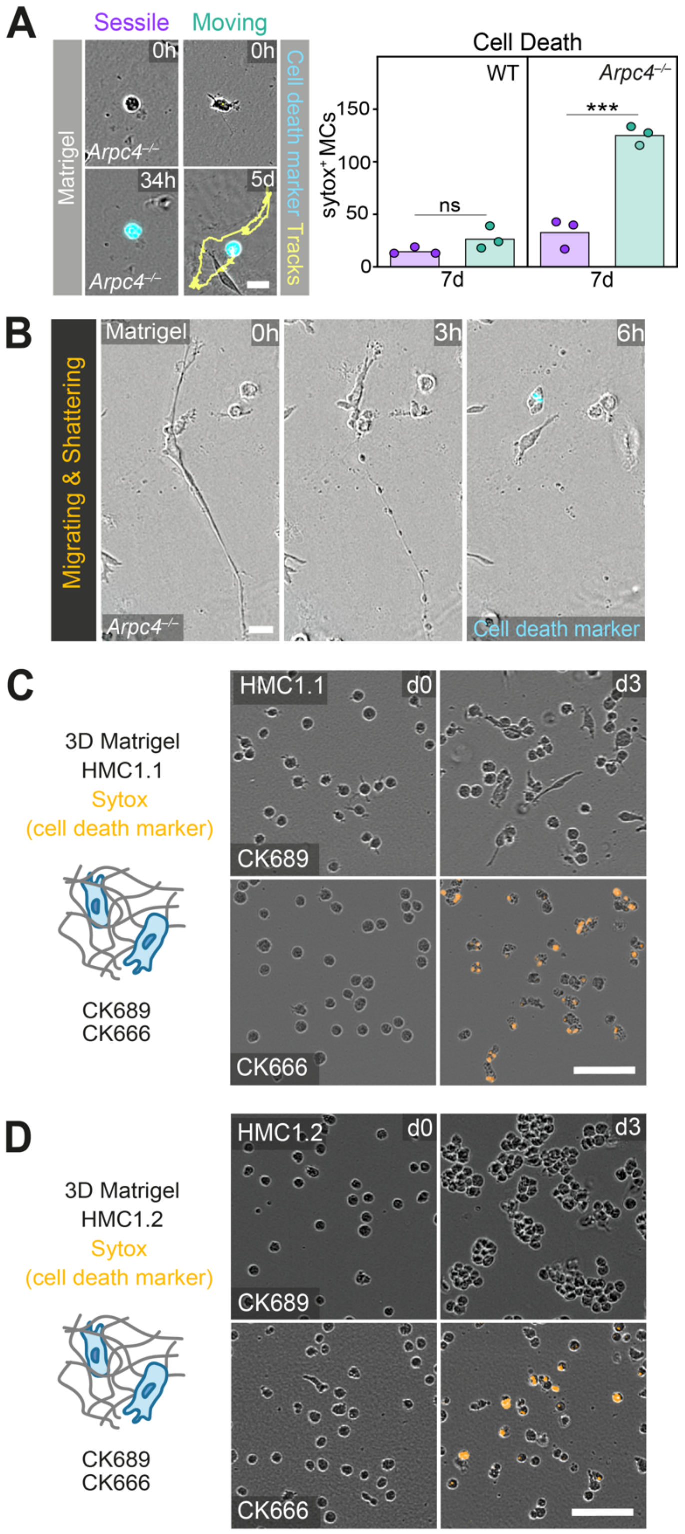


**Fig. S4: Analysis of *Arpc4*$^{-/-}$ BMMC death and functional consequences of Arp2/3 complex inhibition for MC leukemic cell lines.**

**(A)** Potential functional relationship between *Arpc4*$^{-/-}$ BMMC movement and cell death was studied in 3D Matrigel. WT and *Arpc4*$^{-/-}$ BMMCs were observed over 7 d in 3D Matrigel. Sessile and moving MCs were categorized, before the total number of dying cells in each category was quantified. Bars display the mean; ns, non-significant, ***$P \leq 0.001$, *t* test. **(B)** Representative brightfield imaging time series over 6 h of a *Arpc4*$^{-/-}$ BMMC that moves, elongates and shatters in a 3D Matrigel and then immediately undergoes cell death (blue). **(C, D)** Pharmacological inhibition of Arp2/3 complex function and its effects on the survival of human MC lines HMC-1.1 and HMC-1.2 in 3D Matrigel. HMC-1 cells were treated with CK666 or the inactive control CK689. Brightfield images of HMC1.1 and HMC1.2 cells in 3D Matrigel in the presence of the cell death marker Sytox (orange) are displayed (*t*=0 h, *t*=48 h). Scale bars: 30 µm (A), 50 µm (B), 80 µm (C, D).

**Fig. S5.**

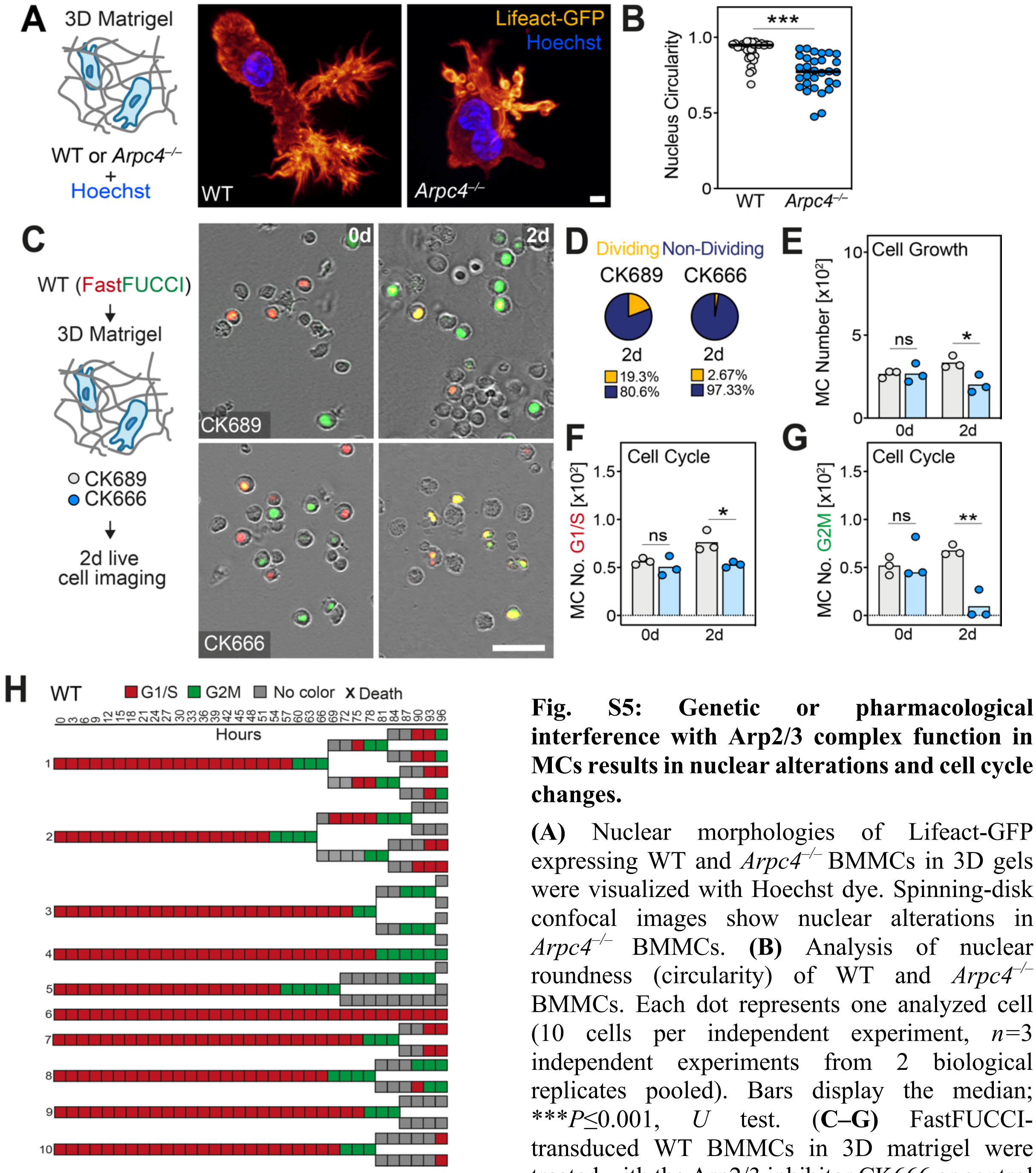


**Fig. S5: Genetic or pharmacological interference with Arp2/3 complex function in MCs results in nuclear alterations and cell cycle changes.**

**(A)** Nuclear morphologies of Lifeact-GFP expressing WT and *Arpc4*$^{-/-}$ BMMCs in 3D gels were visualized with Hoechst dye. Spinning-disk confocal images show nuclear alterations in *Arpc4*$^{-/-}$ BMMCs. **(B)** Analysis of nuclear roundness (circularity) of WT and *Arpc4*$^{-/-}$ BMMCs. Each dot represents one analyzed cell (10 cells per independent experiment, $n$=3 independent experiments from 2 biological replicates pooled). Bars display the median; ***$P \leq 0.001$, *U* test. **(C–G)** FastFUCCI-transduced WT BMMCs in 3D matrigel were treated with the Arp2/3 inhibitor CK666 or control compound CK689. (C) Representative brightfield images are shown ($t$=0 d, $t$=2 d). (D) Quantification of the number of dividing and non-dividing cells over 2 days. The displayed percentages represent average values obtained from $n$=3 independent experiments per genotype. (E–G) Quantification of cell growth and cell cycle stages at day 0 and day 2. Each dot represents one independent experiment ($n$=3 independent experiments per genotype). (E, F) Bars display the mean; ns, non-significant, *$P \leq 0.05$, *t* test. (G) Bars display the median (d0) or the mean (d2); ns, non-significant, U test (d0), **$P \leq 0.01$, *t* test (d2). **(H)** Representative single cell fate mappings of 10 FastFUCCI WT BMMCs over 96 h (in relation to Fig. 4I). Scale bars: 3 µm (A), 80 µm (C).

**Fig. S6.**

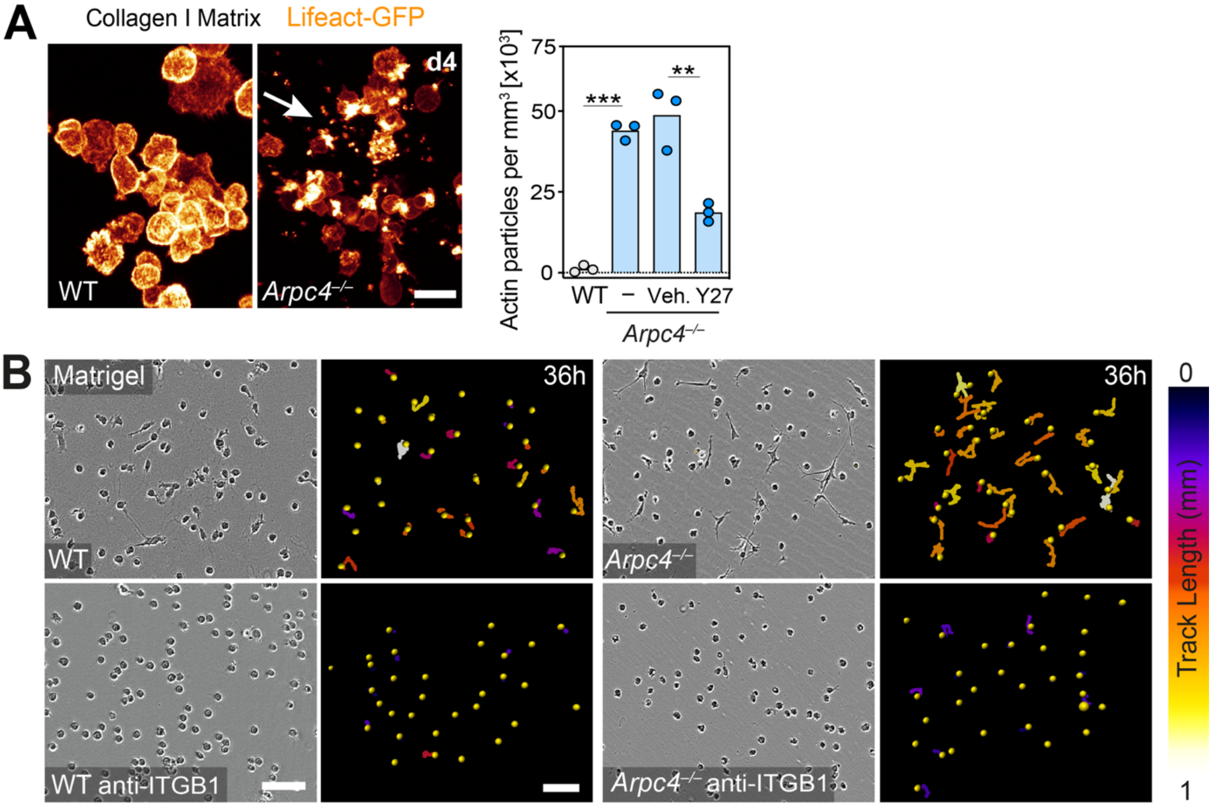


**Fig. S6: Cell fragment formation and anti-ITGB1-mediated rescue of *Arpc4*$^{-/-}$ BMMCs.**

**(A)** Spinning-disk confocal microscopy of *Arpc4*$^{-/-}$ Lifeact-GFP BMMCs show the formation of actin-containing cell fragments (white arrow) in non-adhesive collagen I gels. Since *Arpc4*$^{-/-}$ BMMCs proliferate normally in collagen I gels (Fig. 5C-G), fragment formation is unrelated to the cell cycle arrest and death of *Arpc4*$^{-/-}$ MCs in adhesive 3D Matrigel. Actin fragment numbers were analyzed in a defined volume of collagen I gels and displayed per volume in mm$^3$. The contribution of actomyosin contraction for fragment formation was analyzed by treating *Arpc4*$^{-/-}$ BMMCs with the Y-27632 inhibitor. Each dot represents the average value of three imaging fields of view from one independent experiment ($n$=3 independent experiments from 2 biological replicates). The bars display the mean; ***$P \leq 0.001$, **$P \leq 0.01$, $t$ test. **(B)** The contribution of integrin β1 (ITGB1)-mediated adhesion on BMMC migration in 3D Matrigel was studied. Visualization of WT and *Arpc4*$^{-/-}$ BMMC migration with anti-ITGB1 treatment in 3D Matrigel over 36 h. Representative brightfield images and migration trajectories over 36 h are shown for individual WT and *Arpc4*$^{-/-}$ cells. Anti-ITGB1-treated BMMCs display round cell morphologies and stay rather sessile. Scale bars: 20 µm (A), 80 µm (phase), 200 µm (trajectories) (B).

**Table S1.**

Table S1 lists all mouse strains used in this study.

| Mouse strain | Source | Identifier |
|---|---|---|
| Tg(Arpc4tm1a)Wtsi<br>(brief: $Arpc4^{flox}$) | Metello Innocenti (University of Milan-Bicocca Italy) | MGI: 4433308 |
| Tg(UBC-Brainbow1.0L)35/Orl/J<br>(brief: Ubow) | Marc Bajénoff (CIML, Marseille, France) | MGI: 5645781<br>EMMA: EM 06050 |
| Tg(CAG-EGFP)#Rows<br>(brief: Lifeact-GFP) | Roland Wedlich-Söldner (University Münster, Germany) | MGI: 4831036 |
| Tg(Cma1-cre)ARoer<br>(brief: Mcpt5-Cre) | Axel Roers (TU Dresden, Germany) | MGI: 3785000 |
| Tg(Nckap1l)tm1.2Sixt<br>(brief: $Hem1^{flox}$) | Theresia Stradal (Helmholtz Centre for Infection Research, Braunschweig, Germany) | MGI: 6197558 |
| $Tln1^{tm4.1Crit}$ (brief: $Tln1^{flox}$) | S. Monkley and D. Critchley (University of Leicester, United Kingdom) | MGI: 3785000 |
| $Was^{tm1Kas}$ ($Was^{-/-}$) | Katherine Siminovitch (Mount Sinai Hospital Toronto, Canada) | MGI: 3028420 |
| $Mcpt5\text{-}Cre^{+/-}$ $Arpc4^{fl/fl}$ $Ubow^{+/-}$ | In-house breeding | N/A |
| $Mcpt5\text{-}Cre^{+/-}$ $Arpc4^{fl/fl}$ $Lifeact\text{-}GFP^{+/-}$ | In-house breeding | N/A |
| $Mcpt5\text{-}Cre^{+/-}$ $Arpc4^{fl/fl}$ $Tln1^{fl/fl}$ | In-house breeding | N/A |
| $Mcpt5\text{-}Cre^{+/-}$ $Hem1^{fl/fl}$ | In-house breeding | N/A |

### Table S2.

Table S2 lists the statistical analysis used in each figure. The data was tested for normal distribution with the Shapiro-Wilk test for sample sizes < 10 and with the D´Agostino & Pearson test for sample sizes > 10. Two-tailed unpaired *t* tests and analysis of variance (ANOVA) were performed after data were confirmed to fulfil the criteria of normal distribution and equal variance, otherwise two-tailed Kruskal–Wallis tests or Mann–Whitney *U* tests were applied. If overall ANOVA or Kruskal–Wallis tests were significant, we performed post hoc test with pair-wise comparisons (ANOVA: Tukey, Kruskal–Wallis: Dunn). Stars indicate significance (**P*≤0.05, ***P*≤0.01, ****P*≤0.001). NS indicates non-significant difference (*P*>0.05). Statistical analysis was performed using Prism software (GraphPad Software, Inc,Version 9.3.1).

| **Figure** | **Statist. test** | | **Posthoc test** | **Summary** |
|---|---|---|---|---|
| **1E** | | | | |
| Top | *t* test | *t=2.795, df=58, P=0.0070* | | ** |
| Below | *t* test | t=7.465, df=4 P=0.0017 | | ** |
| **1K** | | | | |
| *Arpc4*$^{-/-}$ - Y27 | Kruskal-Wallis test | P=0.0001 | Dunn´s multiple comparison | *** |
| *Arpc4*$^{-/-}$ - Bleb | Kruskal-Wallis test | P<0.0001 | Dunn´s multiple comparison | *** |
| Y27 | Kruskal-Wallis test | P=0.0005 | Dunn´s multiple comparison | *** |
| Bleb | Kruskal-Wallis test | P<0.0001 | Dunn´s multiple comparison | *** |
| **1M** | | | | |
| WT | Kruskal-Wallis test | P<0.0001 | Dunn´s multiple comparison | *** |
| *Arpc4*$^{-/-}$ | Kruskal-Wallis test | P<0.0001 | Dunn´s multiple comparison | *** |
| **S1E** | Mann-Whitney *U* test | U=738.5, P=0.0003 | | *** |
| **S1F** | *t* test | t=3.669, df=4, P=0.0214 | | * |
| **S1H** | | | | |
| *Arpc4*$^{-/-}$ - Y27 | one-way ANOVA | P<0.0001 | Tukey's multiple comparisons | *** |
| *Arpc4*$^{-/-}$ - Bleb | one-way ANOVA | P<0.0001 | Tukey's multiple comparisons | *** |
| Y27 | one-way ANOVA | P<0.0001 | Tukey's multiple comparisons | *** |
| Bleb | one-way ANOVA | P<0.0001 | Tukey's multiple comparisons | *** |
| **S1K** | | | | |
| WT | one-way ANOVA | P=0.0010 | Tukey's multiple comparisons | ** |
| *Arpc4*$^{-/-}$ | one-way ANOVA | P=0.0002 | Tukey's multiple comparisons | *** |
| **2C** | | | | |
| 3wk | *t* test | t=2.658, df=4, P=0.0565 | | ns |
| 11-13wk | *t* test | t=12.61, df=4, P=0.0002 | | *** |
| 27wk | *t* test | t=29.71, df=4, P<0.0001 | | *** |
| **2E** | *t* test | t=4.576, df=3, P=0.0196 | | * |
| **2G** | *t* test | t=0.7864, df=4, P=0.4756 | | ns |
| **S2C** | | | | |
| *Was*$^{-/-}$ | *t* test | t=0.9902, df=60, P=0.3260 | | ns |
| *Hem1*$^{-/-}$ | *t* test | t=0.8357, df=60, P=0.4067 | | ns |
| **3B** | | | | |
| 0d | *t* test | t=0.1670, df=6, P=0.8729 | | ns |
| 7d | *t* test | t=7.594, df=6, P=0.0003 | | *** |
| **3C** | | | | |

| | | | | |
|---|---|---|---|---|
| 0d<br>7d | *t* test<br>*t* test | t=0.5774, df=6, P=0.5847<br>t=7.531, df=6, P=0.0003 | | ns<br>*** |
| **3D**<br>0d<br>7d | <br>-<br>*t* test | <br>-<br>t=0.3780, df=4, P=0.7247 | | <br>-<br>ns |
| **3E**<br>0d<br>7d | <br>*t* test<br>*t* test | <br>t=0.3152, df=4, P=0.7683<br>t=0.3046, df=4, P=0.7759 | | <br>ns<br>ns |
| **3H**<br>Left<br>Right | <br>*t* test<br>*t* test | <br>t=27.02, df=4, P<0.0001<br>t=18.12, df=4, P<0.0001 | | <br>***<br>*** |
| **3K**<br>Avidin<br>GFP (Ubow) | <br>*t* test<br>*t* test | <br>t=0.2976, df=6, P=0.7760<br>t=4.100, df=6, P=0.0064 | | <br>ns<br>** |
| **3M Growth**<br>0d<br>6d<br>**3M Death**<br>0d<br>6d | <br>*t* test<br>*t* test<br><br>-<br>*t* test | <br>t=0.5288, df=4, P=0.6249<br>t=5.607, df=4, P=0.0050<br><br>-<br>t=4.397, df=4, P=0.0117 | | <br>ns<br>**<br><br>-<br>* |
| **3N Growth**<br>0d<br>3d<br>**3N Death**<br>0d<br>3d | <br>*t* test<br>*t* test<br><br>*t* test<br>*t* test | <br>t=2.163, df=4, P=0.0966<br>t=32.14, df=4, P<0.0001<br><br>t=1.000, df=4, P=0.3739<br>t=30.59, df=4, P<0.0001 | | <br>ns<br>***<br><br>ns<br>*** |
| **3O Growth**<br>0d<br>3d<br>**3O Death**<br>0d<br>3d | <br>*t* test<br>*t* test<br><br>-<br>*t* test | <br>t=0.1930, df=4, P=0.8564<br>t=7.989, df=4, P=0.0013<br><br>-<br>t=7.946, df=4, P=0.0014 | | <br>ns<br>**<br><br>-<br>** |
| **S3C**<br>0d<br>2d | <br>Mann-Whitney *U* test<br>*t* test | <br>U=12, P=0.7000<br>t=9.413, df=4, P=0.0007 | | <br>ns<br>*** |
| **S3D**<br>0d<br>2d | <br>*t* test<br>*t* test | <br>t=1.220, df=4, P=0.2895<br>t=3.667, df=4, P=0.0215 | | <br>ns<br>* |
| **S3E**<br>0d<br>2d | <br>*t* test<br>*t* test | <br>t=1.818, df=4, P=0.1432<br>t=0.5397, df=4, P=0.6180 | | <br>ns<br>ns |
| **S3I** | *t* test | t=4.641, df=4, P=0.0097 | | ** |
| **4D**<br>0d<br>7d | <br>*t* test<br>*t* test | <br>t=2.613, df=4, P=0.0592<br>t=60.21, df=4, P<0.0001 | | <br>ns<br>*** |
| **4E**<br>0d<br>7d | <br>*t* test<br>*t* test | <br>t=2.429, df=4, P=0.0721<br>t=7.654, df=4, P=0.0016 | | <br>ns<br>** |
| **4F**<br>0d<br>7d | <br>-<br>*t* test | <br>-<br>t=22.53, df=4, P<0.0001 | | <br>-<br>*** |
| **4G**<br>G1/S<br>G2M | <br>*t* test<br>*t* test | <br>t=0.7371, df=6, P=0.4889<br>t=0.2618, df=6, P=0.8022 | | <br>ns<br>ns |
| **4N**<br>0d<br>6d | <br>*t* test<br>*t* test | <br>t=0.3106, df=4, P=0.7716<br>t=2.635, df=4, P=0.0579 | | <br>ns<br>ns |
| **4O**<br>0d<br>6d | <br>-<br>*t* test | <br>-<br>t=0.5799, df=4, P=0.5931 | | <br>-<br>ns |

| | | | | |
|---|---|---|---|---|
| **4Q** | | | | |
| 0d | - | - | | - |
| 6d | *t* test | t=0.1527, df=4, P=0.8860 | | ns |
| **4R** | | | | |
| 0d | *t* test | t=1.264, df=4, P=0.2749 | | ns |
| 6d | *t* test | t=0.9459, df=4, P=0.3978 | | ns |
| **S4A** | | | | |
| WT | *t* test | t=1.830, df=4, P=0.1412 | | ns |
| *Arpc4*$^{-/-}$ | *t* test | t=9.485, df=4, P=0.0007 | | *** |
| **5A** | | | | |
| BSA | *t* test | t=0.3004, df=8, P=0.7715 | | ns |
| Col1 | *t* test | t=0.8527, df=8, P=0.4186 | | ns |
| Matrigel | *t* test | t=0.09416, df=8, P=0.9273 | | ns |
| FN | *t* test | t=1.063, df=8, P=0.3190 | | ns |
| Laminin | *t* test | t=0.8801, df=8, P=0.4045 | | ns |
| **5B** | | | | |
| 24h | *t* test | t=2.445, df=4, P=0.0708 | | ns |
| 48h | *t* test | t=7.663, df=4, P=0.0016 | | ** |
| **5D** | | | | |
| 0d | *t* test | t=0.8134, df=4, P=0.4616 | | ns |
| 7d | *t* test | t=0.08501, df=4, P=0.9363 | | ns |
| **5E** | | | | |
| 0d | *t* test | t=0.07293, df=4, P=0.9454 | | ns |
| 7d | *t* test | t=2.447, df=4, P=0.0707 | | ns |
| **5F** | | | | |
| 0d | *t* test | t=0.3586, df=4, P=0.7380 | | ns |
| 7d | *t* test | t=1.265, df=4, P=0.2744 | | ns |
| **5G** | | | | |
| 0d | Mann-Whitney *U* test | U=4, P>0.9999 | | ns |
| 7d | *t* test | t=0.5633, df=4, P=0.6033 | | ns |
| **5I** | | | | |
| 7d WT - WT ITGB1 | one-way ANOVA | P=0.9919 | Tukey's multiple comparisons | ns |
| 7d WT ITGB1 – *Arpc4*$^{-/-}$ ITGB1 | one-way ANOVA | P=0.7202 | Tukey's multiple comparisons | ns |
| 7d *Arpc4*$^{-/-}$ - *Arpc4*$^{-/-}$ ITGB1 | one-way ANOVA | P=0.0099 | Tukey's multiple comparisons | ** |
| **5J** | | | | |
| 7d WT - WT ITGB1 | one-way ANOVA | P=0.6224 | Tukey's multiple comparisons | ns |
| 7d WT ITGB1 – *Arpc4*$^{-/-}$ ITGB1 | one-way ANOVA | P=0.4240 | Tukey's multiple comparisons | ns |
| 7d *Arpc4*$^{-/-}$ - *Arpc4*$^{-/-}$ ITGB1 | one-way ANOVA | P=0.0070 | Tukey's multiple comparisons | ** |
| **S5B** | Mann-Whitney | U=108.5, P<0.0001 | | *** |
| **S5E** | | | | |
| 0d | t test | t=0.04775, df=4, P=0.9642 | | ns |
| 2d | t test | t=3.518, df=4, P=0.0245 | | * |
| **S5F** | | | | |
| 0d | t test | t=0.9048, df=4, P=0.4167 | | ns |
| 2d | t test | t=3.540, df=4, P=0.0240 | | * |
| **S5G** | | | | |
| 0d | Mann-Whitney | U=4, P>0.9999 | | ns |
| 2d | t test | t=6.283, df=4, P=0.0033 | | ** |
| **6B** | | | | |
| WT – *Arpc4* | one-way ANOVA | P=0.0046 | Tukey's multiple comparisons | ** |
| WT – *Arpc4Tln1* | one-way ANOVA | P=0.5814 | Tukey's multiple comparisons | ns |
| *Arpc4* – *Arpc4Tln1* | one-way ANOVA | P=0.0134 | Tukey's multiple comparisons | * |

| | | | | |
|---|---|---|---|---|
| **S6A** | | | | |
| WT – *Arpc4*$^{-/-}$ | t test | t=25.77, df=4, P<0.0001 | | *** |
| Veh. – Y27 | t test | t=5.236, df=4, P=0.0064 | | ** |

### Table S3.

Table S3 lists all antibodies, cell lines and chemicals used in this study.

| Antibody | Company | Identifier |
|---|---|---|
| Rat Anti-Mo CD117 (cKIT) FITC | Thermo Fisher Scientific | Cat# 11-1171-81<br>RRID: AB_465185 |
| Armenian Hamster Alexa Fluor 647 anti-FcεRIα clone: MAR-1 | Biolegend | Cat# 134310<br>RRID: AB_1626093 |
| β-Actin (C4) HRP | Santa Cruz Biotechnology | Cat# sc-47778 HRP<br>RRID: AB_2714189 |
| Donkey pAb to Goat IgG - HRP | Abcam | Cat# ab6885<br>RRID: AB_955423 |
| Donkey Anti-rabbit Alexa Fluor 488 | Thermo Fisher Scientific | Cat# A21206<br>RRID: AB_2535792 |
| Goat Anti-rabbit Alexa Fluor 405-conjugated | Thermo Fisher Scientific | Cat# A-31556;<br>RRID: AB_221605 |
| Goat Anti-GFP Dylight™ 488-conjugated | Rockland | Cat# 600-141-215;<br>RRID: AB_1961516 |
| Goat anti-ARPC4 Antibody | Everest Biotech | Cat# EB08249<br>RRID: AB_2274344 |
| Goat anti-ARPC1A Antibody | Abcam | Cat# ab133160<br>RRID:<br>AB_11154940 |
| Mouse Anti-smooth muscle actin Cy3-conjugated | Sigma-Aldrich-Merck | Cat# C6198;<br>RRID: AB_476856 |
| Rabbit Anti-collagen IV | Abcam | Cat# ab19808;<br>RRID: AB_445160 |
| Purified NA/LE Hamster anti-Rat CD29 | BD Biosciences | Cat# 555002<br>RRID: AB_395636 |
| Rat Anti-CD16/CD32 Antibody | BD Biosciences | Cat# 553142;<br>RRID: AB_394657 |
| Rabbit anti-ARPC2 Antibody | Abcam | Cat# ab133315 |
| Rabbit anti-ARP3 Antibody | Abcam | Cat# ab181164<br>RRID: AB_2892539 |
| Swine Anti-rabbit HRP-conjugated Antibody | Agilent Dako | Cat# P0217<br>RRID: AB_2728719 |
| Anti-human-KIT microbeads CD117 | Miltenyi Biotec | Cat# 130-091-332 |
| anti-FcεRI-FITC | eBiosciene / Thermo Fisher Scientific | Cat# 11-5899-42<br>RRID:<br>AB_10732835 |
| anti-CD117-PE | Miltenyi-Biotec | Cat# 130-111-593<br>RRID: AB_2654579 |
| PE/Dazzle$^{TM}$ 594 anti-mouse CD206 (MMR) Antibody | BioLegend | Cat# 141731<br>RRID: AB_2565932 |
| Alexa Fluor 488 anti-mouse CD206 (MMR) Antibody | BioLegend | Cat# 141710<br>RRID:<br>AB_10933252 |
| **Cell line** | **Company** | **Identifier** |
| CHO (Chinese hamster ovary) cells for SCF production | Provided by Georg Häcker (University of Freiburg, Germany) | N/A |

| | | |
|---|---|---|
| WEHI-3 cells for IL-3 production | Provided by Rudolf Grosschedl (MPI of Immunobiology and Epigenetics, Freiburg, Germany) | CVCL_3622 |
| PlatE cells for Retrovirus production | Provided by Angelika Rambold (MPI of Immunobiology and Epigenetics, Freiburg, Germany) | N/A |
| HMC-1.1 Human Mast Cell Line | Merck | Cat# SCC067 |
| HMC-1.2 Human Mast Cell Line | Merck | Cat# SCC062 |
| **Chemicals** | **Company** | **Identifier** |
| Avidin | Thermo Fisher Scientific | Cat# A2667 |
| Avidin FITC | Sigma-Aldrich-Merck | Cat# A2050-2ML |
| Blebbistatin | Merck | Cat# 203390 |
| CK666 | Merck | Cat# 182515 |
| CK689 | Merck | Cat# 182517 |
| Fibronectin from human plasma | Sigma-Aldrich-Merck | Cat# F2006 |
| Fluoromount G® Mounting Medium | Thermo Fisher Scientific | Cat# 00-4958-02 |
| Hoechst | Thermo Fisher Scientific | Cat# 62249 |
| Laminin | Roche | Cat# 11243217001 |
| Matrigel | Corning | Cat# 354234 |
| Propidium Iodide Staining | Abcam | Cat# ab129817 |
| PureCol (Bovine Collagen I) | Advanced BioMatrix | Cat# 5005-100ml |
| Sytox Green nucleic acid stain | Thermo Fisher Scientific | Cat# S7020 |
| Sytox Orange nuclei acid stain | Thermo Fisher Scientific | Cat# S11368 |
| Y-27632 | Merck | Cat# 688001 |
| Dispase | Corning | Cat# 354235 |
| CLS-1 - Collagenase, Type 1 | CellSystems GmbH | Cat# LS004197 |
| Hyaluronidase | Sigma-Aldrich-Merck | Cat# H3506-5G |
| Human Serum Type AB Male | PAN Biotech | Cat# P30-2901 |
| Amphothericin B | Corning | Cat# 30-003-CF |
| CELL MEM-NEAA sterile, 100x ,10 mM, CELLPURE® | Roth | Cat# 9185.1 |
| Alpha-Monothioglycerol | Sigma-Aldrich-Merck | Cat# M-6145 |
| toluidine blue O | Sigma-Aldrich-Merck | Cat# T-3260 |

| | | |
|---|---|---|
| Tris-(2-carboxyethyl)-phosphin –hydrochlorid solution (TCEP); 0.5M, pH7.0 | Sigma-Aldrich-Merck | #646547 Sulpeco; CAS: 51805-45-9 |
| $MgCl_2$ (Magnesium chloride*hexahydrate, BioXtra) | Sigma-Aldrich-Merck | #M2670 CAS: 7791-18-6 |
| EDTA (Ethylenediaminetetraacetic acid disodium salt dihydrate) | Sigma -Aldrich-Merck | #E5134 CAS: 6381-92-6 |
| Benzonase (nuclease) | EMD Millipore | #70664 CAS: 9025-65-4 |
| Chloroacetamide | Sigma-Aldrich-Merck | #22790 CAS: 79-07-2 |
| Ethanol (absolute, suitable for HPLC, ≥99.8%) | Sigma-Aldrich-Merck | #34852-M CAS: 64-17-5 |
| Ammonium bicarbonate | Honeywell/Fluka | #40867 CAS1066-33-7 |
| RapiGest SF | Waters | #186002123 CAS: 308818-143-5 |
| LysC (Lysyl Endopeptidase) | Wako | #125-02543 CAS: 78642-25-8 |
| SpeedBeads magnetic carboxylate Modified Particles (hydrophobic) | Cytiva | #65152105050250 |
| SpeedBeads magnetic carboxylate Modified Particles (hydrophilic) | Cytiva | #45152105050250 |
| Trypsin Sequencing Grade Modified | Promega | #V5113 |
| Acetonitrile (suitable for HPLC, gradient grade, ≥99.9%) | Sigma-Aldrich-Merck | #34851 CAS: 75-05-8 |
| Water ($H_2O$) LC-MS Grade | Pierce | #85189 CAS: 7732-18-5 |
| Trifluoroacetic acid | Pierce | #28903 CAS: 76-05-01 |
| Formic acid | Fisher Scientific | #A11750 CAS: 64-18-6 |
| ReproSil-Pur 120 C18-AQ, 1.9 µm reversed phase chromatography beads | Dr. Maisch Laboratories (72119 Ammerbuch-Entringen, Germany) | #r119.aq. |
| CoAnn Empty Self-pack NanoLC Column Tube with Emitter Tip and without frit, 360 µm OD x 75 µm ID x 8 µm Tip x 50cm L | MSWIL (5735 GW, Aarle-Rixtel, Netherlands) | #ICT36007508-50-5 |

**Legends to Movies:**

**Movie S1.**

**Migration of WT and *Arpc4*$^{-/-}$ BMMCs in 3D Matrigel.**

MCs were cultivated from WT and *Arpc4*$^{\Delta MC}$ mice and applied to a 3D Matrigel assay in the presence of SCF. Cell movement occurred in the 3D network of the solidified Matrigel. Phase contrast brightfield microscopy over 36 h revealed faster movement of *Arpc4*$^{-/-}$ BMMCs with more elongated morphologies and needle-like protrusions at cellular leading edges. This video relates to Figs. 1, C to E and figs. S1, E and F.

**Movie S2.**

**Lobopodia-like protrusion dynamics of *Arpc4*$^{-/-}$ BMMCs in 3D Matrigel.**

MCs were cultivated from *Arpc4*$^{\Delta MC}$ *Lifeact-GFP*$^{+/-}$ and WT *Lifeact-GFP*$^{+/-}$ mice and applied to a 3D Matrigel assay in the presence of SCF. Video sequences on top display maximum intensity projections, videos below display a z-plane of a zoomed-in cell extension at the leading edge of a WT or *Arpc4*$^{-/-}$ BMMC. Spinning-disk confocal microscopy revealed that WT BMMCs form F-actin rich, small lamellopodial-like extensions at the leading edge.
In contrast, *Arpc4*$^{-/-}$ BMMCs form lobopodia-like, cylindrical cell protrusions covered with bleb-like structures. This video relates to Figs. 1, F and G.

**Movie S3.**

***Arpc4*$^{-/-}$ BMMCs lose cell fragments from cellular edges and the cell body in 3D Matrigel.**

MCs were cultivated from *Arpc4*$^{\Delta MC}$ *Lifeact-GFP*$^{+/-}$ mice and applied to a 3D Matrigel assay in the presence of SCF. The first video sequence displays the max. projection of a *Arpc4*$^{-/-}$ BMMC over 45 min. The second videos display a maximum intensity projection of a *Arpc4*$^{-/-}$ BMMC (left) and a zoom-in of the same cell's cell body (right) over 2 min. Spinning-disk confocal microscopy revealed that *Arpc4*$^{-/-}$ BMMCs lose actin-containing cellular fragments from cell edges and the cell body (indicated with white arrows). This video relates to Fig. 1H and fig. S1G.

**Movie S4.**

***Arpc4*$^{-/-}$ BMMCs undergo cell death in 3D Matrigel over seven days.**

MCs were cultivated from WT and *Arpc4*$^{\Delta MC}$ mice and applied to a 3D Matrigel assay in the presence of SCF and a cell death marker (orange). BMMCs were observed with phase contrast brightfield microscopy over seven days (168 h). The video shows that *Arpc4*$^{-/-}$ BMMCs display deficient proliferation in 3D Matrigel, with a gradually increasing cell death rate over time. This video relates to Figs. 3, A, B and D.

**Movie S5.**

**Cell cycle visualization of WT and *Arpc4*$^{-/-}$ BMMCs in 3D Matrigel.**

MCs were cultivated from WT and *Arpc4*$^{\Delta MC}$ mice and transduced with FastFUCCI to visualize the cell cycle stage by color: G1/S phase (red nucleus), G2/M phase (green nucleus). BMMCs were applied to a 3D Matrigel assay in the presence of SCF and imaged with phase contrast brightfield microscopy over seven days (168 h). The first video sequence displays a representative WT BMMC over 17.5 h that transitions from the G1/S phase to the G2/M phase and then divides. The second video sequence displays an overview of transduced WT BMMCs over four days (96 h). Several WT cells undergo cell division (highlighted with white arrows), distributing throughout the Matrigel. The third video sequence displays an overview of transduced *Arpc4*$^{-/-}$ BMMCs over four days. Most *Arpc4*$^{-/-}$ BMMCs fail to transit from the G1/S phase into the G2/M phase and several cells undergo cell death while being stuck in G1/S phase. The fourth video sequence displays the different cell fates of *Arpc4*$^{-/-}$ BMMCs in 3D Matrigel over four days while visualizing the cell cycle dynamics with the FastFUCCI construct. This video relates to Figs. 4, A to K.

**Movie S6.**
**Blocking integrin-mediated mechano-coupling rescues *Arpc4*$^{-/-}$ BMMC proliferation.**
*Arpc4*$^{-/-}$ BMMCs were treated with an anti-ITGB1 antibody to block the integrin β1-mediated adhesion to ECM components in 3D Matrigel. The video shows a representative overview of *Arpc4*$^{-/-}$ BMMCs, either untreated or treated with anti-ITGB1, in the presence of a cell death marker (orange). Cells were observed with phase contrast brightfield microscopy over 28 h. *Arpc4*$^{-/-}$ BMMCs with the anti-ITGB1 treatment display a round cell morphology, stay rather immotile and some cells undergo cell divisions (highlighted by white arrows) in the adhesive 3D Matrigel. This video relates to Figs. 5, H to J and fig. S6B.
